# Calcium signalling is required for anterior patterning in the mouse embryo

**DOI:** 10.1101/2024.11.14.623160

**Authors:** Matthew J. Stower, Richard C. V. Tyser, Shifaan Thowfeequ, Felix Zhou, Marta Portela, Konstantinos Miti, Jacintha Sugnaseelan, Xin Lu, Shankar Srinivas

## Abstract

Anterior-Posterior axis formation in the mouse embryo requires the active migration of the DVE cell population at E5.5. While intracellular Ca^2+^ signalling has been shown to control cell migration in multiple cell contexts, it is unknown whether it is required for DVE migration. The pattern of Ca^2+^ activity in the mouse embryo at early peri-implantation stages is also unknown. Using the GCaMP6f Ca^2+^ reporter line we performed a detailed assessment of Ca^2+^ dynamics between E0.5 – E5.5 using live imaging. We find that prior to implantation, Ca^2+^ transients are rare, but at E5.5 widespread, periodic, Ca^2+^ transients in extra-embryonic tissues can be observed, including in the VE and ExE. In contrast, cells of the E5.5 epiblast remain relatively quiescent but show sporadic large-scale multi-cellular waves. Inhibition of SERCA at E5.5 abolishes Ca^2+^ transients and leads to DVE arrest, indicative that these transients are required for axial patterning. Together these results reveal the pattern of Ca^2+^ handling in the early mouse embryo and a novel requirement in anterior-posterior axis formation.

## INTRODUCTION

The distal visceral endoderm (DVE) is a specialised cell population in the embryonic day 5.5 (E5.5) mouse embryo that undergoes a collective migration required for anterior-posterior axis specification (Thomas, Brown et al. 1998, Srinivas, Rodriguez et al. 2004, Stower and Srinivas 2018). DVE cells, induced at the distal tip of the E5.5 egg-cylinder, migrate over the course of 3-5 hours (Srinivas, Rodriguez et al. 2004) towards one-side of the embryo where they secrete inhibitors of the TGF-β/NODAL (Yamamoto, Saijoh et al. 2004) and WNT (Belo, Bouwmeester et al. 1997) pathways that are required for primitive streak formation. The primitive streak is the site of gastrulation and beings to form from ∼E6.5 on the side of the epiblast farthest from the inhibitory signals secreted by the DVE (Ding, Yang et al. 1998, Tam and Loebel 2007). Mutants where the DVE is not induced, or in which DVE migration is aberrant or inhibited lead to incorrectly patterned, non-viable embryos (Ding, Yang et al. 1998, Thomas, Brown et al. 1998, Huelsken, Vogel et al. 2000, Brennan, Lu et al. 2001).

DVE cells are a migratory epithelial cell population that retain intact tight- and adherent-junctions, defined by ZO-1 and E-cadherin (Trichas, Joyce et al. 2011), throughout their migration. However, they also show characteristics traditionally associated with mesenchymal cells – highly dynamic basally located projections polarised in the direction of migration (Srinivas, Rodriguez et al. 2004, Migeotte, Omelchenko et al. 2010). As DVE cells remain within the monolayer epithelium of the visceral endoderm (VE), they must actively regulate both apical and basal domains throughout migration.

Regulation of intracellular calcium (Ca^2+^) signalling is involved in a diverse array of cellular responses including: fertilisation cell migration, proliferation, apoptosis, and differentiation (Clapham 2007, Paudel, Sindelar et al. 2018, Stewart and Davis 2019, Sukumaran, Nascimento Da Conceicao et al. 2021). Cells can generate a transient cytosolic increase in Ca^2+^ through mobilisation from intracellular stores (including the endoplasmic reticulum (ER), Golgi apparatus and mitochondria), or from extracellular sources (Clapham 2007). This is controlled by ion pumps and channels including the sarcoplasmic/endoplasmic reticulum Ca^2+^ ATPase (SERCA)(Sanderson, Charles et al. 1994). Cells can also signal cell-to-cell via gap junctions (Lin, Rurangirwa et al. 2004), causing a sequential wave of Ca^2+^ transients that can span multiple cells in a tissue (Leybaert and Sanderson 2012). Intracellular Ca^2+^ can regulate migration by influencing the cytoskeleton, cell polarity and focal adhesions (Tsai, Kuo et al. 2015). For example, in mesenchymal cells, Ca^2+^ can form a front-rear (high-low) gradient, polarising cells to form lamellipodia only at the front of the cell while mediating rear detachment of protrusions through regulation of focal adhesions (Tsai, Kuo et al. 2015, Kim, Lee et al. 2016). In some cases such as in *Xenopus* axial mesoderm, no intracellular gradient is observable but Ca^2+^ is elevated in the leading cells of the migratory cell population (Hayashi, Yamamoto et al. 2018).

In the mouse oocyte, sperm entry triggers a transient cytosolic release of Ca^2+^ within 3 minutes, followed by a series of Ca^2+^ oscillations every 20-30 min over the course of several hours (Cuthbertson, Whittingham et al. 1981, Lawrence, Whitaker et al. 1997, Deguchi, Shirakawa et al. 2000, Stewart and Davis 2019). This is an event highly conserved across vertebrate taxa, though species differ in the number and frequency of subsequent oscillations (Whitaker 2006)(Mizuno, Sassa et al. 2013) (Wozniak, Mayfield et al. 2017). In the mouse 2-cell stage, it has been shown that small vesicles located at the sub-plasmalemmel cytoplasm, termed Membrane Associated RNA-containing Vesicles (MARVs) have a higher level of Ca^2+^ than the cytosol (Wang, Yasmin et al. 2022) and persist until embryonic day 3.5 (E3.5) (Wang, Yasmin et al. 2022). MARVs act as a critical Ca^2+^ reservoir for the regulation of mitochondrial activity, and transfer Ca^2+^ from membrane located MARVs to the mitochondria that surround the nucleus at these early stages, though no cytoplasm-wide Ca^2+^ transients have been reported (Wang, Yasmin et al. 2022).

After this stage, our knowledge of Ca^2+^ signalling in early vertebrate development comes mainly from studies in fish, amphibians and chick (Ferrari and Spitzer 1999, Wallingford, Ewald et al. 2001, Creton 2004, Kreiling, Balantac et al. 2008, Papanayotou, De Almeida et al. 2013, Eno, Gomez et al. 2018, Hayashi, Yamamoto et al. 2018, Lee, Oliveira et al. 2024). Ca^2+^ oscillations have only been observed in the mouse node at E7.5 (Takao, Nemoto et al. 2013) and the forming cardiac crescent from E7.75 (Tyser, Miranda et al. 2016), constituting evidence for continued Ca^2+^ transients during early mouse development (Stewart and Davis 2019). Therefore, there exists a gap in our knowledge of any role of Ca^2+^ in the mouse at early stages.

It has been possible to visualise Ca^2+^ transients during early development using fluorescent Ca^2+^ indicators such as f-aequorin (Leclerc, Webb et al. 2000), Calcium Green-1 dextran (Wallingford, Ewald et al. 2001) and Fluo-4 AM (Wang, Yasmin et al. 2022). However, there are technical difficulties to these approaches in the mouse embryo, particularly as it grows, including the dilution of injected reporters as cells divide, and the necessity for diffusion of the dye throughout the depth of tissues. To circumvent these issues we made use of a genetically encoded fluorescent Ca^2+^ reporter mouse line GCaMP6f (Chen, Wardill et al. 2013, Madisen, Garner et al. 2015) to perform a detailed stage-series assessment of Ca^2+^ dynamics between E0.5 - E5.5 using time-lapse microscopy. We find that at up to E3.5 Ca^2+^ oscillations are rare, stochastic events, while at E4.5 there is a higher level of Ca^2+^ in the trophectoderm and primitive endoderm compared to the epiblast. At E5.5 there is a significant up-regulation in the frequency and amplitude of oscillations mainly in the extra-embryonic tissues of the extraembryonic ectoderm (ExE and VE). The epiblast remains relatively quiescent, but we observe occasional multi-cellular Ca^2+^ waves that spread across this tissue. Inhibition of SERCA abolishes Ca^2+^ transients at E5.5 and causes the arrest of DVE migration, an event required for anterior-posterior axis formation.

## METHODS

### Mouse strains

B6;129S-*Gt(ROSA)26Sor*^*tm95*.*1(CAG-GCaMP6f)Hze*^/J (Madisen, Garner et al. 2015) that contain a Cre-inducible GCaMp6f reporter were crossed to B6.Cg-*Edil3*^*Tg(Sox2-cre)1Amc/*^J, Sox2-Cre (Hayashi, Lewis et al. 2002) mice to generate mice that constitutively express the GCaMP6f reporter. Offspring were bred to subsequently segregate out the Sox2-Cre and establish a constitutive, ubiquitous GCaMP6f reporter line. GCaMP6f homozygous stud males were crossed with *Gt(ROSA)26Sor*^*tm4(ACTB-tdTomato,-GFP)Luo*^ mT/mG (Muzumdar, Tasic et al. 2007) or CD1 (Charles River) females to generate embryos for experiments. For long-term culture experiments, we used the Hex-GFP mouse line (Rodriguez, Casey et al. 2001) where DVE cells are marked by cytoplasmic EGFP. Homozygous Hex-GFP stud males (with and without the mT/mG allele) were mated with CD1 females (Charles River). All mice were maintained on a 12 h light, 12 h dark cycle. Noon on the day of finding a vaginal plug (0.5 days *post coitum*) was designated embryonic day (E)0.5. Embryos of the appropriate stage were dissected in M2 medium (Sigma) with fine forceps and tungsten needles as previously described (Srinivas 2010)

### Confocal Microscopy – Live Imaging

Dissected embryos were transferred to a 8-well Lab-Tek II chambered cover-glass slide (Sigma) with 250 µl of culture medium (1:1 knock-out serum replacement: CMRL, supplemented with 1% L-glutamine) pre-equilibrated at 37°C and 5% CO_2_. The imaging slide was mounted into a pre-warmed stage and chamber of an inverted, ZIESS LSM 880 confocal microscope. Embryos were imaged at a range of durations and intervals (**see Table S1**). For more detailed analysis of Ca^2+^ dynamics in E5.5 embryos a single mid-sagittal z-section was imaged at an interval of 5 seconds for a duration of 10 minutes using a water immersion 40x/1.2 NA objective. Each developmental stage comprised a freshly dissected litter.

### Lightsheet Microscopy

For lightsheet imaging experiments, 20 µl glass capillaries (Brand, 701904) with plungers were used to create a 2% agarose cylinder, in which a copper wire with a diameters of 150 um (for E5.5 experiments) was inserted. Once the agarose solidified, the wire was removed to leave a hollow lumen into which an embryo could be carefully inserted. The glass capillary was then mounted into the imaging chamber of a Zeiss Z1 lightsheet microscope and the region of the agarose cylinder containing the mounted embryo extruded into an imaging chamber filled with pre-heated culture medium (as above). Imaging was performed using a 63×/1.0 NA Plan-Apochromat water immersion objective with dual-side illumination. Z-stack volumes were then captured at a 10 second interval for 10 minutes (see Table S1). Extended focus projections were generated by ZEN software (ZEISS).

### Lattice Lightsheet Microscopy (LLSM)

For LLSM, a ZEISS Lattice Lightsheet 7 microscope equipped with a 13.3x 0.4 NA illumination objective, 44.83x 1.0 NA water immersion detection objective with an Alvarez manipulator and a pre-defined Sinc3 100µm x 1800nm light sheet beam was used. The heating and CO_2_ concentrations were maintained at 37°C and 5% respectively using the IncuControl system (Version 1.0.3; Ibidi). Embryos were stabilised between two glass rods within the chamber of a 8-chambered #1.5 µ-slide (Ibidi) filled with culture medium (as above), so that the lightsheet scans through the embryo along its anterior–posterior axis. X-stack volumes were obtained at 0.8 µm x-intervals at imaging intervals of 2 seconds for 20 minutes (to visualise intracellular Ca^2+^ signal propagation), 20 seconds for 5 hours (to image Ca^2+^ waves) or 5 minutes for 6 hours (for DVE migration live imaging). The resulting image stacks were deskewed with coverglass correction using the ZEN Blue Pro software (Version 3.10, ZEISS).

### Pharmacological inhibitor experiments

GCaMP6f embryos from one or more litter were randomly separated into wells of an 8-well Lab-Tek II chambered cover-glass slide filled with culture medium (as above). For short-term inhibitor experiments, each embryo was imaged at a single mid-sagittal z-level for 2 minute duration, with a 10 second interval between images on a pre-heated, climate controlled chamber (37°C 5% CO_2_) of a LSM 880 (ZEISS) confocal microscope. An initial time-lapse represented a baseline number of Ca^2+^ transients for each embryo (the sum of all transients in all tissues in the time-lapse). Next the selected concentration of inhibitor (10nM thapsigargin or 200nM cyclopiazonic acid, CPA) or control solvent (DMSO) was added to each well, and embryos were cultured for 30 mins. Embryos were then re-imaged and the number of Ca^2+^ transients per embryo calculated. Data for each inhibitor comprised a minimum of 3 experimental repeats with multiple embryos in each experiment. For CPA experiments an additional recovery time-point was acquired after washing off the inhibitor and culturing for an additional 30 minutes in fresh culture medium. Embryos were then re-imaged and the total number of Ca^2+^ transients calculated. For long-term culture experiments homozygous Hex-GFP stud males were crossed with CD1 females and embryos dissection at E5.5. Embryos from one or more litter were randomly separated into wells of an 8-well Lab-Tek II chambered cover-glass slide filled with culture medium (as above) and imaged on an LSM 880 (ZEISS) confocal microscope. Images of each well were taken before 10nM thapsigargin or DMSO was added. Embryos were then cultured for 8 hours, then re-imaged and the number of embryos that migrated was scored. Experiments comprised data from a total of 5 independent experiments.

### Automated Analysis of Ca^2+^ transients in E5.5 embryos

Supervised machine learning using a convolutional neural network (CNN) classifier was used to detect the peaks of cytosolic Ca^2+^ transients in the GCaMP6f time-lapse data of E5.5 embryos. To train the CNN, a training dataset was created by manually identifying the time-point of the peak of Ca^2+^ transients in three E5.5 time-lapse datasets. These time-points were used to construct an idealised probability Ca^2+^ imaging trace where the intensity is non-zero only at the peak time-point. Given an input Ca^2+^ imaging trace, and its idealised equivalent a CNN was trained to predict the probability of observing a Ca^2+^ transient peak within a symmetrical temporal window of N=35 time-points. All raw GCaMP6f time-lapse data (Ca^2+^ trace) were first de-trended using asymmetric least squares fitting (Eilers and Boelens 2005) prior to CNN input. Unbiased clustering of Ca^2+^ transients per embryo was carried out to assess the similarity of Ca^2+^ dynamics in each embryo in an unbiased manner. Individual Ca^2+^ traces were pairwise compared to construct a similarity matrix with similarity was defined as the largest positive value of the signal cross-correlation. A hierarchical density based clustering method (HDBSCAN), was then applied using the Python HDBSCAN package (Campello, Moulavi et al. 2013) to the similarity matrix to cluster Ca^2+^ traces. Global clustering across all embryos was performed by computing the similarity matrix pooling together all Ca^2+^ traces from all embryos. Individual traces have different minimum and maximum intensities within individual embryos. Traces were first normalised per embryo by z-score, using the mean and standard deviation of all traces from the respective embryo. HDBSCAN generates the optimal clustering number given min_cluster_size – the minimum number of samples in a group for that group to be considered a cluster and min_samples – the number of samples in a neighbourhood for a sample to be considered as a core point. The output clusters include ‘core’ clusters, whereby traces have within a ‘core’ cluster have high intra-similarity and a ‘outlier’ cluster, which contain traces determined insufficiently similar to a trace with core clusters, and insufficient intra-similarity to form a new core cluster. This approach compares similarity based on the shape of Ca^2+^ traces and is therefore tolerant to slight misalignment in peak times, and peak-to-peak distances. For the global clustering, qualitative examination of the outlier cluster showed traces had a similar spiking pattern but were asynchronous in time. Therefore we also used the ‘outlier’ class as a cluster in the global clustering result. We label the clusters numerically, in ascending order of the mean transient number of traces in the cluster. Therefore in our global clustering, cluster 2 is also the outlier class (Fig 3C). As quantitative measures of Ca^2+^ peak behaviour, we computed the mean number of peaks, mean intensity of peaks, periodicity as the mean peak-to-peak distance, and the mean duration of peaks within a trace. Peak duration was measured per peak as the difference between start and end peak times. We define peak start time as the timepoint just before half peak intensity and end time as the timepoint just after half peak intensity.

### Kymograph visualisation of GCaMP6f Ca^2+^ reporter activity

To visualise GCaMP6f Ca^2+^ reporter activity in time-lapses of E5.5 embryos, individual traces were stacked together as a kymograph, each row being the normalised intensity trace, each column a time-point, with signal intensity coloured using the ‘magma’ colour-scheme. The yellower/whiter the colour, the stronger the trace intensity. The purpler/ blacker the colouring, the weaker the trace intensity. Traces were normalised by min-max normalisation, (trace – min)/(max – min) where min and max are the minimum and maximum intensity across all traces within an embryo. The colour-scheme was then applied to the value range [0,1]. Traces are row-sorted by performing an indirect stable sort with cluster number first and the time of its first peak second. For global clustering, where individual traces were z-scored per embryo instead of min-max normalised, the same row-sorting is applied but with the kymograph colouring applied to the positive value range [0, 2].

### Spatiotemporal analysis of Ca^2+^ transients in E5.5 embryos

To investigate the spatiotemporal pattern of calcium transients in E5.5 GCaMP6f:membrane-tdTomato time-lapses obtained at 5 second interval for 10 minutes duration we used Motion Sensing Superpixel (MOSES) tracking (Zhou, Ruiz-Puig et al. 2019). As Ca^2+^ transients occurring sequentially in neighbour cells or from region to region within a cells, will appear to move like a wave, we used super pixel motion tracking to analyse the behaviour of the GCaMP6f reporter. We applied MOSES with Farnebäck optical flow (Farnebäck 2003) to extract the mean temporal Ca^2+^ flow pattern for each embryo. To ensure minimal bleed-through of cell movements, a binary threshold of mean + 0.5 standard deviation of the calcium signal was used prior to tracking. To enable comparison across embryos, all embryos were rotated to be vertical with ExE top by applying principal components analysis to the cell centroids. Flow maps showing the average directional motion behaviour of the calcium transients were then generated. For flow maps of epiblast waves, we applied MOSES with TV-L1 optical flow (Pock, Urschler et al. 2007, Wedel, Pock et al. 2009) which is more suitable for capturing large displacements.

### Whole-mount Immunofluorescence

Whole-mount immunofluorescence was performed as previously described (Trichas, Joyce et al. 2011). Briefly, embryos were fixed in 4% PFA for 20 minutes at room temperature then washed three times for 5 minutes each in 0.1% Triton-X100 in phosphate buffered saline (PBS); incubated in 0.25% Triton-X100 in PBS for 25 minutes; washed three times in 0.1% Tween-20 in 1xPBS; then blocked with 2.5% donkey serum, 2.5% goat serum, and 3% Bovine Serum Albumin (BSA) in 0.1% Triton-X100 in PBS overnight. Embryos were then incubated at 4°C overnight in primary antibodies diluted in 1:100 in block. Embryos were washed three times in 0.1% Tween-20 in PBS (PBT) for 5 minutes each, with a final wash of 15 minutes; incubated overnight at 4°C with appropriate secondary antibody 1:100 in 0.1% PBT and 1nM phalloidin overnight at 4°C. Finally embryos were washed four times for 5 minutes in PBT at room temperature; and mounted with Vectashield mounting media containing 4′,6-diamidino-2-phenylindole (DAPI) (Vector Labs H-1200

### Antibodies and stains

Primary antibodies used were: rabbit anti-Cleaved Caspase-3 (Cell Signalling, 9961S, Lot: 43 (06/2014)), rabbit anti-Phospho-Histone H3 (S28) (Cell Signalling, 9713P, Lot: 2 (09/2013)), rabbit anti-OCT-4 (Abcam, ab200834, Lot: GR212680-16), rabbit anti-CDX2 (Cell Signalling, 9775, Lot: 3(12/2017)). All primary antibodies were used at 1:100 dilution. Secondary antibodies used were: donkey anti-rabbit Alexa Fluor 555 (Invitrogen, A31570, Lot: 1945911). For F-actin staining phalloidin-Atto 647N (Sigma, 65906) was used at a 1nM final concentration in 1 x PBT. Nuclei were stained by Vectashield mounting media containing 4′,6-diamidino-2-phenylindole (DAPI) (Vector Labs H-1200).

### Confocal microscopy of fixed samples

Fixed E5.5 embryos were imaged following the immunofluorescence protocol (as above) on a LSM 880 (ZEISS) confocal microscope using a 40x oil (1.36NA) objective. Z-stacks of embryos were acquired at 1 µm interval using non-saturating parameters. Max intensity projections (MIPs) were made using ZEN black software (ZEISS). Figures were prepared with Adobe Photoshop and Adobe Illustrator (Adobe Inc.).

### Analysis of apoptosis in thapsigargin treated embryos

For analysis of apoptosis embryos the level of Cleaved-Caspse-3 (Casp3) was compared between thaspigargin treated and DMSO as a negative control. 3D z-stacks of cultured embryos stained for F-actin and Casp3 were used to analysed the percentage of staining in each embryo as follows; first the F-actin channel was used to manually segment the 3D volume of each embryo using FIJI. The Casp3 channel was then automatically segmented using FIJI Particle Analyzer plugin, enabling the percentage of Casp3 staining in each embryo to be calculated. Statistical tests were carried out using R (ver 1.1.45).

## RESULTS

### Ca^2+^ transients in the pre-implantation mouse embryo are rare stochastic events

In order to image cytosolic Ca^2+^ transients in all tissues of the early mouse embryo, we generated a constitutively and ubiquitously expressed version of the GCaMP6f Ca^2+^ reporter line (Madisen, Garner et al. 2015), henceforth referred to as ‘GCaMP6f’ (see Methods). We crossed this line with *Gt(ROSA)26Sor*^*tm4(ACTB-tdTomato,-GFP)Luo*^ *‘*mT/mG’ mice (Muzumdar, Tasic et al. 2007) to generate embryos in which cells and tissues could be discerned with a fluorescent membrane-tdTomato signal while recording cytosolic Ca^2+^ dynamics with the GCaMP6f reporter. To investigate whether cytosolic Ca ^2+^ transients occur at early developmental stages, we therefore performed a stage-series of high-time resolution confocal time-lapse imaging of freshly dissected embryos from a range of stages (E0.5, E1.5, E2.5, E.3.0, E3.5, E4.5 and E5.5). As Ca^2+^ levels could change slowly or rapidly, in order to minimise photo-toxicity we employed an imaging strategy to accommodate these possibilities using a combination of high time-resolution short-duration (typically 5 second intervals for 10 minutes), and lower time-resolution longer-duration imaging parameters (for example 20 minute intervals for 10 hours) varying with the stage (see Methods and Table S1).

We defined a Ca^2+^ transient as a cytosolic increase in GCaMP6f fluorescence, followed by return to basal levels. If a cell showed more than one Ca^2+^ transient within our imaging duration we termed this a Ca^2+^ oscillation. If Ca^2+^ transients occurred sequentially across neighbouring cells in a directional manner we termed this a wave.

At E0.5 we imaged GCaMP6f:membrane-tdTomato embryos from post-pro-nucleus formation until the 2-cell stage (N=5) (Movie S1). No Ca^2+^ transients were observed in any embryo (N=5). Next, we imaged E1.5 embryos (N=13) from the 2-cell until the 4-cell stage (Movie S2). Though we did not observe any large changes in cytosolic Ca^2+^ concentration, upon division from 2 to 4 blastomeres, a higher relative level of Ca^2+^ appeared along the interface between dividing cells (Movie S2). We also noted that there were multiple punctate regions within the cytosol with a higher relative level of Ca^2+^ (Fig. A-A’, Movie S3) that appear stable over short-term imaging (5-second interval for 10 minutes) (Movie S4). At E2.5 the punctate regions of high GCaMP6f signal were now less obvious around the nuclei, but seemed to be more prevalent near the apical (outer) surface of blastomeres (Fig. 1B-B’, Movie S5). At E3.0 we could still observe cytosolic clusters of puncta similar to those at E2.5 (N=7) (Fig. 1C-C’, Movie S6) and in one embryo, we also observed 3 blastomeres with a single Ca^2+^ transient occurring in our 10 minute imaging duration.

**Figure 1.**
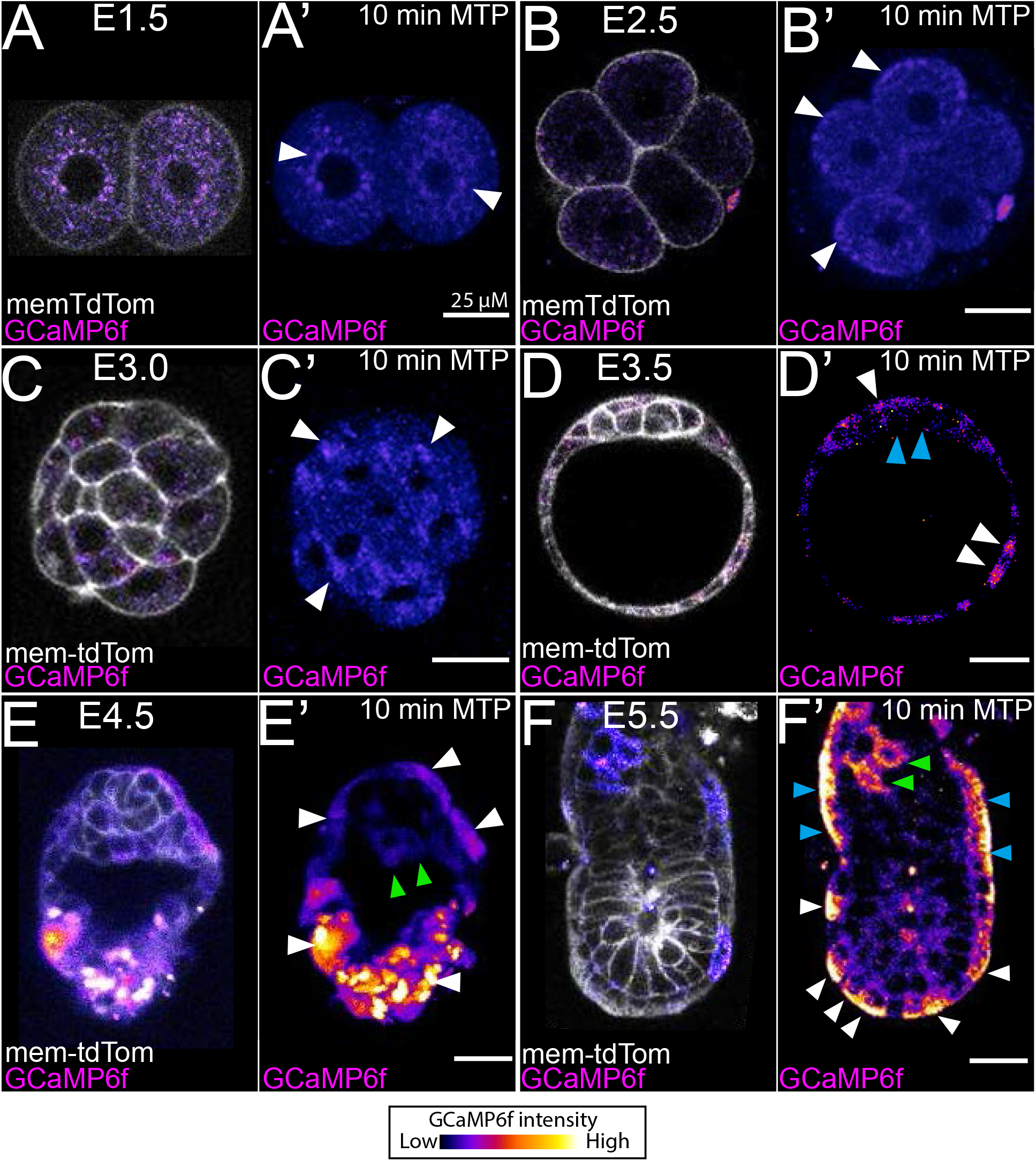
Cytosolic Ca^2+^ oscillations occur at post-implantation stages in the mouse embryo. Representative single z-sections of a single-time point, and 10 minute max-intensity time projections (MTP) from time-lapse imaging of the Ca^2+^ reporter mouse GCaMP6f crossed with membrane-tdTomato at; (A-A’) E1.5 (B-B’), E2.5 (C-C’), E3.0 (D-D’), E3.5 (E-E’), E4.75 and (F-F’). **(A-A’)** E5.5. At E1.5 embryos show puncta with a high GCaMP6 signal around the nucleus (A’, arrowhead). **(B-B’)** E2.5 show puncta with a high level of Ca^2+^ at the apical domain relative to the rest of the cell (arrowheads in B’). **(C-C’)** E2.5 embryos show large groups of puncta with a high GCaMP6 signal but no cytosolic transients (arrowheads in C’). **(D-D’)** At E3.5 trophectoderm cells show a higher level of Ca^2+^ (white arrowheads) than the ICM (blue arrowheads). **(E-E’)** E4.5 show higher level of Ca^2+^ in the trophectoderm (white arrowheads in E’) and primitive endoderm (green arrowheads in E’) than the epiblast. **(F-F’)** At E5.5 widespread Ca^2+^ oscillations with high intensity occur in the emVE (white arrowheads in F’) exVE (blue arrowheads in F’) and ExE (green arrowheads in F’). Each stage represented a fresh dissection. Additional time-lapse imaging durations and intervals were also tested (see Table S1 and Movies S1-9). All scale bars = 25 µM.

Interestingly at E3.5 all embryos (N=6) showed higher levels of Ca^2+^ in trophectoderm cells relative to the ICM (Fig. 1D-D’, Movie S7). A single Ca^2+^ transient was also observed in a trophectoderm cell in one embryo (Movie S7). At E4.5, trophectoderm and primitive endoderm cells showed an elevated level of Ca^2+^ compared to the pluripotent epiblast (N=5) (Fig. 1E-E’, Movie S8), but no Ca^2+^ transients were observed. Together, these findings suggest that between E0.5 - E4.5 Ca^2+^ transients are rare events. At early stages, small cellular vesicles with Ca^2+^ levels higher than the surrounding cytosol are localised either around the nucleus or near the apical domain. From blastocyst stages, extra-embryonic tissues show a relatively higher cytosolic Ca^2+^ level than the pluripotent ICM (E3.5) or epiblast (E4.5).

### E5.5 marks the onset of widespread periodic Ca^2+^ transients

At E5.5, following implantation, the embryo elongates to form the egg-cylinder, with the epiblast abutting the proximally located extra-embryonic ectoderm (ExE), both surrounded by the monolayer epithelium of the visceral endoderm (VE) (termed emVE when overlying the epiblast, and exVE overlying the ExE) (Fig. 2A, see diagram). Strikingly, when imaged at 5-second intervals, we observe periodic Ca^2+^ transients throughout the emVE, exVE (Fig. 2A-A’, Movies S9 and S10), as well as in the ExE (Fig. 2B-B’, Movie S10, top right), with several cells undergoing multiple transients (Ca^2+^ oscillations) within a 10 minute imaging period (N=14). Transients were not temporally coordinated across the embryo, but we noted that they often occurred sequentially in small groups of neighbouring cells in the emVE (Fig. 2D-D’), exVE (Fig. 2D-D’) and ExE (Fig. 2B,D’’, and Movie S10). At this stage a sub-set of VE cells, the DVE, are induced at the distal tip of the egg cylinder and become more columnar than surrounding cells (Fig. 2A-C, green region). Though we could observe Ca^2+^ oscillations in these cells, they did not appear to show Ca^2+^ behaviour distinct from surrounding emVE cells (Fig. 2A-C, see green region). To test whether there was any spatial bias to these events, we applied motion sensing superpixel (MOSES) (Zhou, Ruiz-Puig et al. 2019) analysis (see methods) to show flow patterns of transients across the ExE and VE (Fig. S3). This revealed that there was no directional bias in the transients, which could move either distally (see asterisk in Fig. S3A’’) or proximally (see asterix in Fig. S3B’’) within distinct groups of cells.

**Figure 2.**
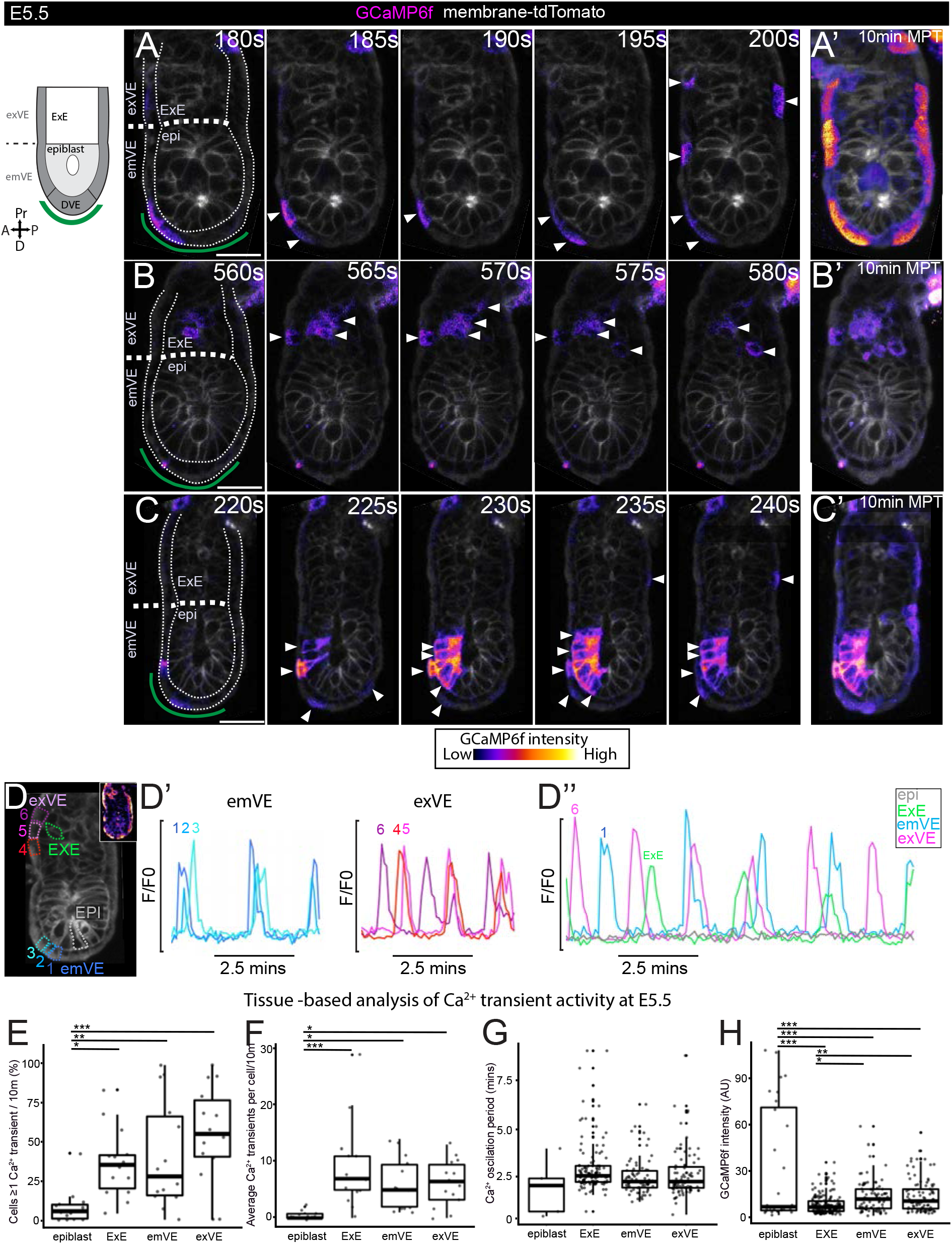
At E5.5 Ca^2+^ transients are more widespread and occur with higher frequency in extra-embryonic than embryonic tissues. **(A-C)** Selected time-points of three E5.5 GCaMP6f:membrane-tdTomato embryos imaged every 5-seconds for 10 minutes showing Ca^2+^ transients in VE, ExE and epiblast. Green line in first frame indicates approx. position of DVE cells based on morphology - no obvious difference in dynamics could be observed in these cells when compared to the surrounding emVE and exVE. (A’, B’, C’) Max-intensity temporal projections (MTP) of full 10 minute time-lapse (MTPs) (see Movie S10**). (D)** E5.5 embryo with ROI’s relating to D-D’’. **(D’-D’’)** Traces of GCaMP6f fluorescence for emVE, exVE showing neighbouring cells undergoing Ca^2+^ oscillations at a similar periodicity.**(D’’)** The timing of Ca^2+^ transients is different across cells. Epiblast cells cells are largely quiescent. **(E-H)** E5.5 embryos (N=14) analysed using a trained CNN peak detector. **(E)** Graph showing percentage of cells in each embryo that showed ≥1 transient. Epiblast have significantly fewer transients than other tissues (one-way ANOVA, p=<0.05, followed by Tukey’s HSD Test on epiblast vs: ExE, p=<0.05, emVE p=<0.01, exVE p=<0.001)). **(F)** Graph of average number of Ca^2+^ transients per cell. Epiblast cells show the fewest transients (one-way ANOVA, p=<0.001, followed by Tukey’s HSD Test on epiblast vs: ExE, p=<0.001, emVE p=<0.05, exVE p=<0.05)). **(G)** Graph of Ca^2+^ oscillation periodicity for all cells. **(H)** Graph showingGCaMP6f fluorescent intensity at Ca^2+^ transient peaks. Epiblast Ca^2+^ transient peak intensity were on average significantly higher than other tissues (one-way ANOVA, p=<0.001, followed by Tukey’s HSD Test on epiblast vs: ExE, p=<0.001, emVE p=<0.001, exVE p=<0.001)). emVE and exVE showed a slightly higher intensity than the ExE (one-way ANOVA, p=<0.001, followed by Tukey’s HSD Test on ExE: vs: emVE, p=<0.05, exVE p=<0.01)). Note that the emVE was not subdivided into DVE and non-DVE for tissue-based analysis. All scale bars = 25 µM.

As this analysis was based on single-optical sections, where a wave travelling out of the imaging plane would be missed, we next captured 3D-volumes of E5.5 embryos at 10-second intervals using a Z.1. lightsheet microscope (Movie S11). This confirmed that though there are Ca^2+^ transients across the VE and ExE, there are no larger wave-like events across these tissues (N=3). Together this suggest local interconnectivity between neighbouring cells exists, but no directional bias.

At this stage Ca^2+^ transients could also be observed in the epiblast, though much less frequently (Fig. 2C and Movie S10, bottom left). In contrast to the VE and ExE, epiblast transients spread progressively across multiple cells in a wave-like manner and appeared to have a higher intensity (Fig. 2C, D’’’ and Movie S10). Unlike at earlier stages (E1.5 - E3.0) no vesicles with a higher level of Ca^2+^ were observed in any cells at E5.5 embryo.

In order to characterise the number, duration and location of Ca^2+^ transients in more detail, we used the membrane-tdTomato signal and manually outlined regions of interest (ROI) around each cell to create a data-set of 1126 cells (323 epiblast, 430 ExE, 205 emVE and 168 exVE from all the E5.5 embryos (N=14) imaged at 5-second interval for 10 minutes. As we lacked a DVE marker we did not sub-divide the emVE into the DVE for this analysis. We manually annotated the peaks of Ca^2+^ transients from three embryos and used this to train a convolutional neural network (CNN) “Ca^2+^ peak detector” (see Methods). We then applied the trained CNN Ca^2+^ peak detector to identify transients in the remaining embryos and analysed at the tissue level: the proportion of cells in each tissue that showed at least one Ca^2+^ transient; the number of Ca^2+^ transients in each cell; the average time between Ca^2+^ transients in a cell (oscillation period); the duration of Ca^2+^ transients and; the relative level of level of GCaMPf fluorescence at Ca^2+^ transient peaks.

We found that the proportion of cells with ≥ 1 Ca^2+^ transient at E5.5 was significantly higher in all extra-embryonic tissues (ExE, emVE, exVE) than the epiblast (Fig. 2E). Only 10% (32/323) of epiblast cells imaged showed ≥ 1 Ca^2+^ transient compared to 38% (163/430) of ExE, 40% (83/205) of emVE and 56% (94/168) of exVE cells (Fig. 2E). The average number of transients per cell was also significantly higher in extra-embryonic (∼5 – 8 transients in 10 minutes) than than epiblast cells (∼1 transient in 10 minutes) (Fig. 2F, see also Table S2). The duration of Ca^2+^ transients (see Methods) was similar across tissues at around 20 seconds (Table S1, include graph?). The average Ca^2+^ oscillation period was similar between cells in extra-embryonic tissues (∼3 minutes), but cells also showed a wide-range of oscillations periods (30 seconds to 9 minutes) (Fig. 2G, Table S2). While the average oscillation period for epiblast was shorter (∼2 minutes) this only represented a total of 5 events across the entire dataset, since an oscillation in the epiblast was a rare event (Fig. 2G). Lastly, we analysed the relative intensity of Ca^2+^ transients. This revealed that epiblast had a greater range cytosolic Ca^2+^ levels compared to the extra-embryonic tissues (Fig. 2H).

### Unbiased clustering of E5.5 cells reveals different cells-types with distinct Ca^2+^ handling behaviours

While these analyses enable comparison of the average behaviour of cells in each tissue, they do not take into account the spatio-temporal pattern of behaviour. We therefore performed an unbiased clustering based analysis of our data using HDBSCAN (Campello, Moulavi et al. 2013), a hierarchical density clustering approach (Fig. 3A-B’, Fig. S1 and see Methods). We also plotted Ca^2+^ dynamics in each embryo as a kymograph to visualise the temporal behaviour of cells across all tissues during 10 minute time-lapse experiments to be easily visualised (Fig. 3B’’, Fig. S2 and see Methods). Using both, we clustered all cells (N=1126) from our E5.5 time-lapse dataset (N=14 embryos) according to the similarity of their Ca^2+^ dynamics and plotted this as an integrated kymograph (Fig. 3C). We found that cells clustered into four groups (clusters 0, 1, 2, 3), with cluster 0 representing cells with no or trace Ca^2+^ transient activity (Fig. 3C, Table S3) over the 10 minute duration of imaging. We then analysed the contribution of cells from tissues to each cluster (Fig. 3D) and plotted this onto spatial maps for each embryo (Fig. 3D’, Fig. S2). This showed that while the epiblast is made up predominantly of cells from c0 (no transient), extra-embryonic tissues are made up of cells from all clusters. Cluster c3 is made up predominantly of ExE, while c2 is made up predominantly of ExE, emVE and exVE.

**Figure 3.**
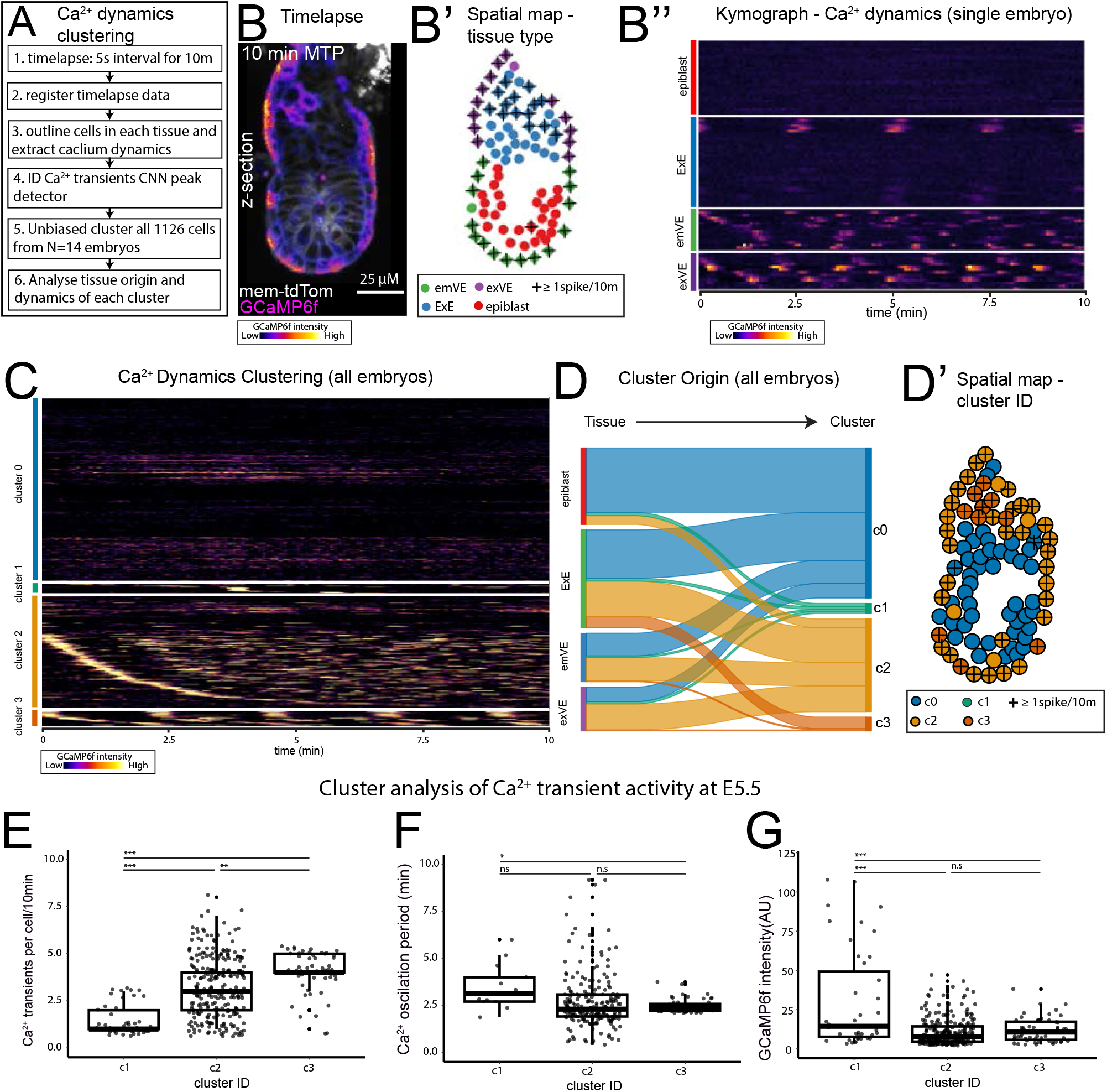
Unbiased clustering of Ca^2+^ dynamics reveals four activity types. **(A)** To assess the similarity of Ca^2+^ transients in all cells from E5.5 (N=14) embryos imaged at a 5-second interval for 10 minutes unbiased clustering of Ca^2+^ transients using a hierarchical density based clustering was performed. **(B)** Example max-intensity time projection of an example E5.5 GCaMP6f embryo. **(B’)** Spatial map of cell-type and Ca^2+^ activity on embryo in B. **(B’’)** Kymograph of E5.5 embryo in B, showing GCaMP6f activity during 10 minutes. **(C)** Kymograph of GCaMP6f activity of 1126 cells from N=14, E5.5 embryos clustered using hierarchical density based clustering. Four clusters were identified with cluster 0 representing little-no activity. **(D)** Sankey graph showing contribution of cells from each tissue to the four cluster types. Epiblast contribute predominantly to c0 (no spike), while extra-embryonic cells contribute to all clusters, more cells from ExE contribute to c3, while the majority of the emVE and exVE contribute to c2. **(D’)** Spatial map of embryo in B, showing cluster ID. **(E-F)**. Quantification of GCaMP6f activity in clusters 1-3. **(E)** Cluster1 had the lowest number of Ca^2+^ transients, while cluster 3 had the highest (one-way ANOVA, p=<0.05, followed by Tukey’s HSD Test on c1 vs: c2, p=<0.001, c3. c2 vs c3:0.01)). **(F)** Ca^2+^ oscillation period showed that cluster 1 had slower oscillations than cluster3 (though c1 data reflected a total of only 13 cells), while there was no signifiant difference between clusters 2 and 3 (one-way ANOVA, p=<0.05, followed by Tukey’s HSD Test on c1 vs: c3, p=<0.05, c2 vs c3: p=>0.05)). **(G)** Intensity of Ca^2+^ transients (AU = arbitrary unit) in each cluster. Cluster 1 transients had on average significantly higher intensity ((one-way ANOVA, p=<0.05, followed by Tukey’s HSD Test on c1 vs: c2, p=<0.001, c3 p=<0.001)).

To differentiate the behaviour of Ca^2+^ transients between the clusters that showed at least one transient (clusters 1 - 3) we analysed the number of Ca^2+^ transients, the oscillation periodicity and intensity of Ca^2+^ transients in each cluster (Fig. 3E-G). This showed that cluster 1 - 3 represent a gradient of lower to higher activity with cluster 1 cells showing fewer transients with the majority of cells showing only 1 transient in 10 minutes (Fig. 3E), a similar oscillation periodicity (Fig. 3F) but at a higher intensity (Fig. 3G) than other clusters, while clusters 2 and 3 show higher numbers of transients, but a similar oscillation periodicity and transient intensity (Fig. 3E-G). Together these results show that cells in the E5.5 embryo can be categorised into different groups based on Ca^2+^ dynamics and that these groups broadly correspond to tissue differences. While epiblast cells are largely confined to either no transients (cluster 0), or infrequent high intensity transients (cluster1), the extra-embryonic tissues largely cluster to higher frequency, average intensity transients (cluster 2-3).

### Ca^2+^ transients in epiblast cells propagate from the apical to basal domain

While Ca^2+^ transients in the epiblast were rare in a single optical section imaged over a 10 minute duration (Fig. 3C and D), when observed, these events were expansive, propagating across multiple cells as a wave (eg: Fig. 2C). To investigate this phenomenon in more detail and across the 3-dimensional extent of the embryo, we imaged GCaMP6f:mTmG embryos using lattice lightsheet microscopy (LLSM), which enabled us to capture events in the near-full volume of the E5.5 embryo and over a longer imaging duration (5 or 20 second interval between volumes over 20 minutes, or 20 second interval for 5 hours, see Table S1). This confirmed our previous observations that epiblast Ca^2+^ transients occurred in a wave-like manner, sequentially propagating across neighbouring cells of the epiblast. Moreover, volume imaging showed that there is variability in the extent of the epiblast across which the wave travelled. In some cases it covering the majority of epiblast cells (Fig. 4A and Movie S12), while in other cases, it was more limited in extent (Fig 4B and Movie S13). We observed a single epiblast wave in 6/13 embryos imaged for 20 minutes (3/10 in embryos imaged at 20-second intervals and 3/3 in those imaged at 5-second intervals). When imaged over a longer, 5 hour duration, 3/3 embryos showed multiple epiblast waves; 2 embryos had 4 waves, 1 embryos had 3 waves. These waves lasted for an average of 70 seconds ±37s, but could be as short as 20 seconds. The time between epiblast waves was on average 25 minutes ±16 minutes, with a maximum interval of 57 minutes.

**Figure 4.**
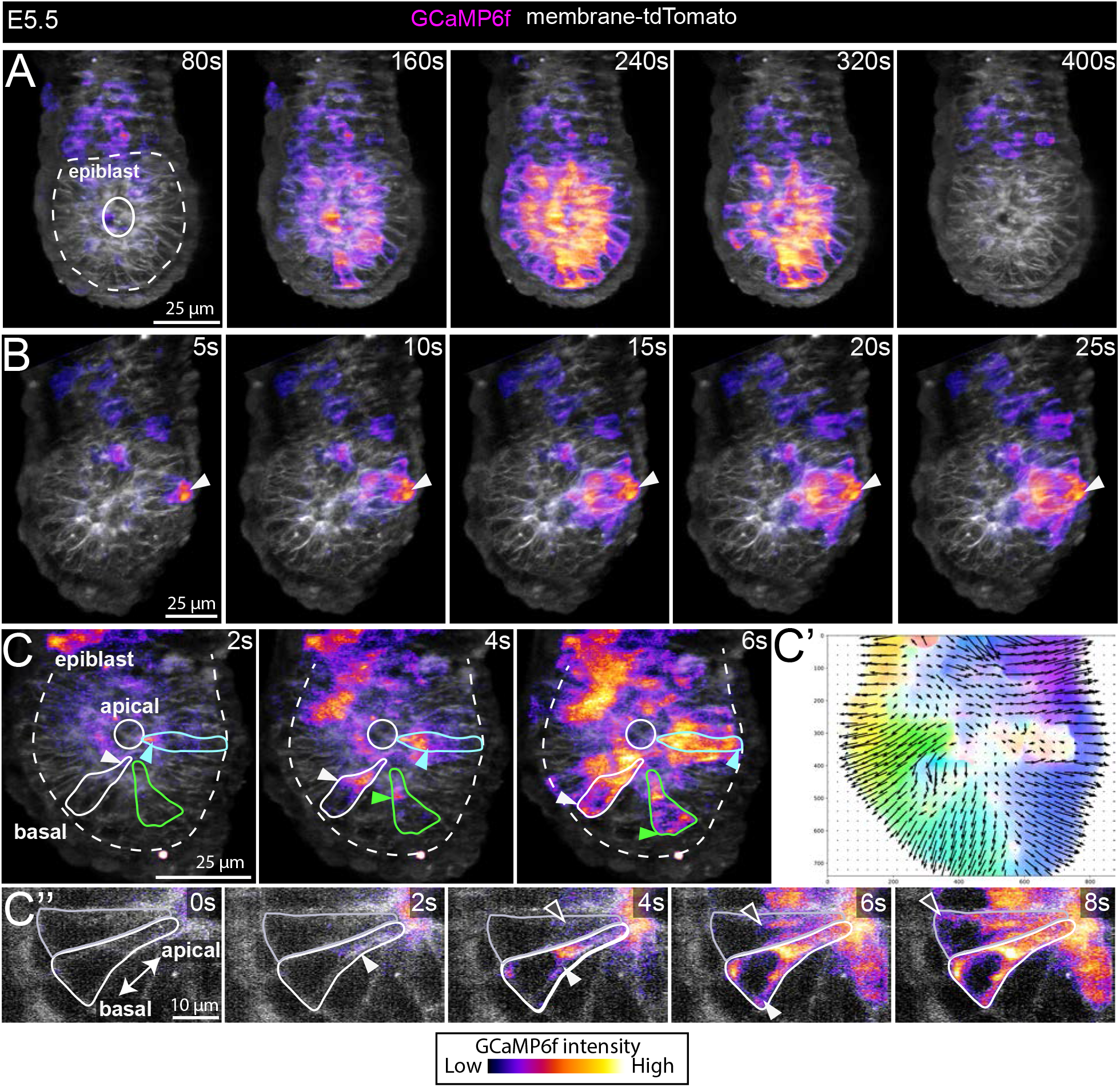
Epiblast Ca^2+^ waves propagate from the apical to basal domain. **(A-C’’)** Selected time-points from movies of E5.5 GCaMP6f embryos imaged using a Lattice lightsheet microscope. **(A)** An example of an epiblast wave involving a large proportion of the tissue captured by imaging a volume of the embryo at 20-second intervals (also see Movie S12). A total of 9/16 embryos showed at least one epiblast Ca^2+^ wave. **(B)** Faster imaging at 5 second interval revealed that the Ca^2+^ wave in the epiblast can be preceded by a transient in a neighbouring VE cell – this was observed in 5/19 events from 9 embryos (white arrowheads) (also see Movie S13). **(C)** Imaging at 2-second interval captured the intracellular propagation of the Ca^2+^ transient; Ca^2+^ activity begins at the apical domain of epiblast cells and moves toward the basal aspect of cells (white-, cyan-, and green-arrowheads denote the position of Ca^2+^ activity in 3 outlined epiblast cells) (N=3 embryos) (see Movie S14). **(C’)** Unbiased super-pixel motion analysis showing the intracellular apical to basal movement in the embryo shown in C. **(C’’)** Selected time-points from a single optical section from the embryo in C showing the apical-basal movement of Ca^2+^ activity in 2 outlined epiblast cells. All images are maximum intensity projections.

In 5/19 waves from the 9 embryos that showed at least one wave, we also noted that the initiation of the epiblast Ca^2+^ wave was preceded by a transient in a VE cell immediately adjacent to the origin of the epiblast wave (Fig. 4B and Movie S13). The Ca^2+^ transient in these VE cells were of high intensity and persisted for longer than the transients in the epiblast cells, returning to resting levels only after the epiblast wave had ended (Fig. 4B and Movie S13). Due to the imaging interval and the fact that we are unable to capture the full embryo volume, it is unclear whether a Ca^2+^ transient in a VE cell always proceeds an epiblast wave, or if this represents only a proportion of these events.

To understand the intracellular propagation of the Ca^2+^ transient within individual cells we imaged embryo volumes at a higher temporal resolution, at 2-second intervals using LLSM. This revealed that Ca^2+^ transients originates from the apical aspect of epiblast cells’ and moves towards their basal aspect (all epiblast cells in N=3 embryos) (Fig. 4C and Movie S14). We confirmed this in an unbiased manner by applying motion sensing super-pixel analysis (MOSES) (Zhou et al., 2019) to track the Ca^2+^ signal as pixel motion (Fig. 4C’).

### Ca^2+^ transients in the E5.5 embryo are dependent on SERCA

Having identified the onset of cytosolic Ca^2+^ transients at E5.5 we next wanted to understand the source of Ca^2+^ these may represent. The endoplasmic reticulum (ER) is a key intracellular Ca^2+^ store. After the release of Ca2+ from these stores into the the cytosol, the sarcoplasmic/ endoplasmic reticulum Ca^2+^-ATPases (SERCA) transport Ca^2+^ from the cytosol back into the ER (Sanderson, Charles et al. 1994) to replenish stores in readiness for the next transient. To determine if the transients we observed were dependent on this mechanism, we cultured GCaMP6f embryos in thapsigargin, a cell-permeable SERCA inhibitor (Thastrup, Cullen et al. 1990, Lytton, Westlin et al. 1991) widely used in developmental studies (Wallingford, Ewald et al. 2001, Creton 2004, Kreiling, Balantac et al. 2008, Schneider, Houston et al. 2008, Bower, Lansdale et al. 2017, Mizuno, Shiozawa et al. 2020). We assessed the total number of Ca^2+^ transients in embryos immediately post-dissection (‘baseline’), and again after 30 minutes of culture with 10nM thapsigargin, or DMSO as a negative control (Fig 5A). Though control embryos cultured in DMSO did show a statistically significant decrease in the number of transients (from an average of 17 at baseline, to 12 Ca^2+^ transients after culture, N=12 embryos), treatment with 10nM thapsigargin resulted in a near complete abolition of Ca^2+^ transients (from an average of 18, at baseline, to 1.3 Ca^2+^ transients post-culture, N=12 embryos). Of these 6/12 had complete loss of Ca^2+^ transients (Fig. 5A-A’’, B).

**Figure 5.**
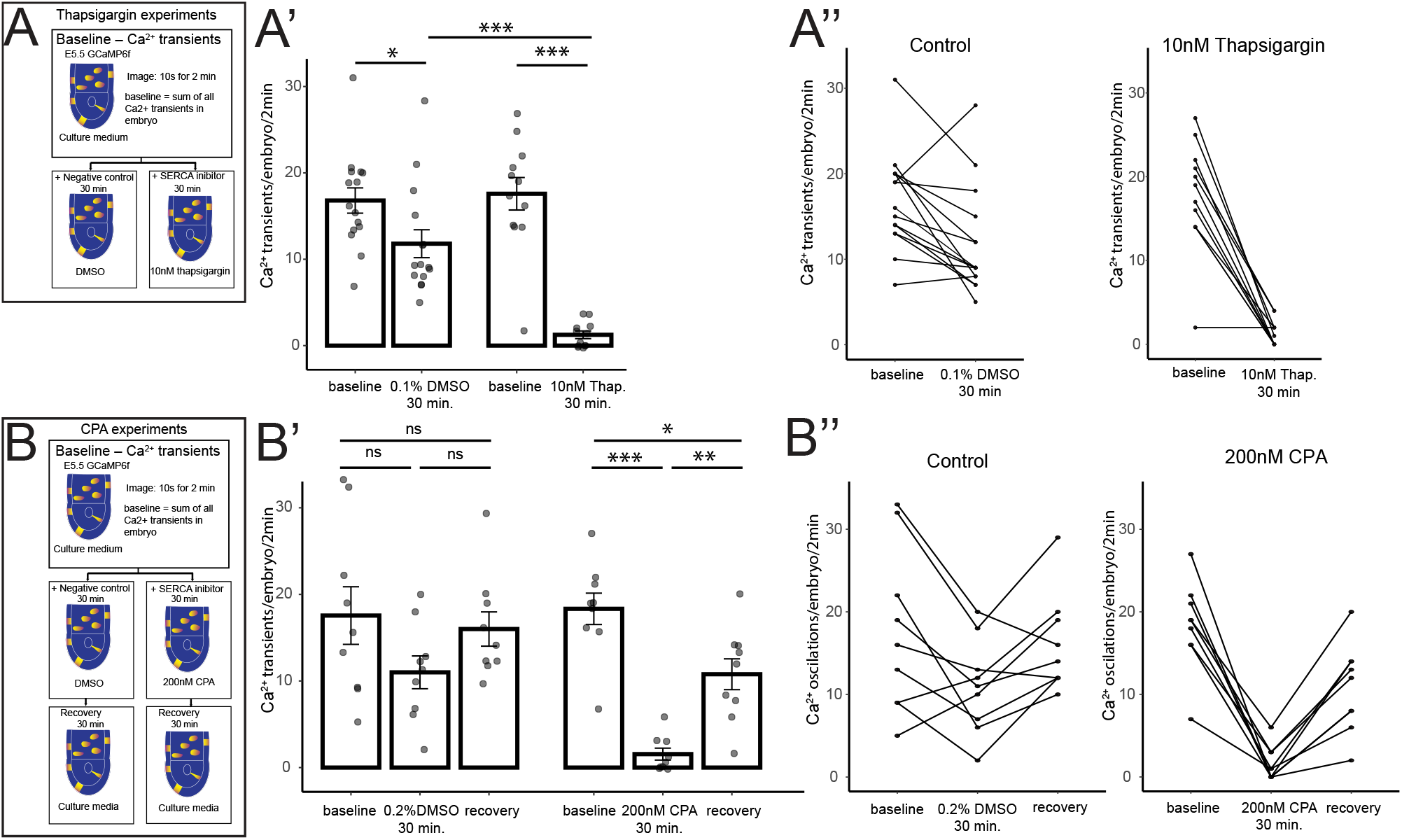
SERCA is required for Ca^2+^ transients at E5.5. To test the requirement of endoplasmic reticulum (ER) for Ca^2+^ dynamics at E5.5, GCaMP6f embryos were treated with SERCA inhibitors and the total number of Ca^2+^ transients measured. **(A)** Diagram of experimental method for non-reversible SERCA inhibitor thapsigargin. Embryos were imaged at a 10-second interval for 2 minutes. **(A’)** Graph of Ca^2+^ transients in embryos treated with 0.1% DMSO and 10nM thapsigargin. DMSO treated embryos had a significant decrease in transients (one-way ANOVA, p=<0.001, followed by Tukey’s HSD Test; baseline vs 30 min DMSO p>0.05), but thapsigargin almost completely abolished transients to a significant effect (one-way ANOVA, p=<0.001, followed by Tukey’s HSD Test, baseline vs 30 min 10nM thapsigargin p=<0.001), that was significantly lower than DMSO controls (one-way ANOVA, p=<0.001, followed by Tukey’s HSD Test, 30 minute DMSO vs 30 min 10nM thapsigargin: p=<0.001). **(A’’)** Graphs of individual embryos from A’, traced from baseline to post-30 minutes culture. **(B)** Diagram of experimental method for the reversible SERCA inhibitor cyclopiazonic acid (CPA). **(B’)** Graph of Ca^2+^ transients in DMSO and 200nM CPA treatment. While 0.2% DMSO treated embryos showed a decrease in transients, it was not significant (one-way ANOVA, p=<0.001, followed by Tukey’s HSD Test, baseline vs 30 minute 10nM thapsigargin: p=>0.05). 200nM CPA treatment almost completely abolished the Ca^2+^ transients to a significant effect (one-way ANOVA, p=<0.001, followed by Tukey’s HSD Test, baseline vs 30 minute 200nM CPA: p=<0.001) but embryos were able to recover significantly after CPA was washed off and cultured with fresh medium (one-way ANOVA, p=<0.001, followed by Tukey’s HSD Test, 30 minute 200nM CPA vs recovery: p=<0.01). **(B’’)** Graphs of individual embryos from B’, traced through-out the experiment.

To further test whether SERCA is required for cytosolic Ca^2+^ activity at E5.5, we treated embryos with a second inhibitor; cyclopiazonic acid (CPA) (Seidler, Jona et al. 1989), also previously used in developmental studies (Creton 2004, Kreiling, Balantac et al. 2008, Bower, Lansdale et al. 2017).

Like thaspigargin, CPA inhibits SERCA but has the advantage that it acts in a reversible manner (Seidler, Jona et al. 1989). We therefore assessed the number of Ca^2+^ transients in E5.5 embryos immediately post-dissection (‘baseline’), after 30 minutes culture with 200nM CPA, or DMSO as a negative control, and then assessed embryos ability to recover after culturing in fresh medium for an additional 30 minutes (Fig. 5C). While DMSO control embryos showed a similar reduction in the number of Ca^2+^ transients as before, though in this case not statistically significant (Fig. 5C-C’’), CPA almost completely abolished all Ca^2+^ transients after 30 minutes culture (from an average of 18, to 1.6 Ca^2+^ transients post-culture, N=9 embryos). Of these 4/9 showed a complete loss of Ca^2+^ transients. (Fig. 5C-C’’, D). CPA treated embryos were able to recover significantly, to approximately half of their initial baseline level after CPA was washed off (**Fig. 5C-C’’, D**). Together these data indicate that Ca^2+^ transient activity at E5.5 is SERCA-dependent.

### Ca^2+^ signalling is required for DVE migration

At E5.5 the DVE cell population undergoes a cell migration within the VE epithelium (Srinivas, Rodriguez et al. 2004). As Ca2+ signalling has been shown to regulate cell migration in a range of cellular contexts (Tsai, Kuo et al. 2015, Kim, Lee et al. 2016, Hayashi, Yamamoto et al. 2018), and given the wide-spread Ca2+ activity in this tissue, we asked whether it is required for the migration of DVE cells. To test whether Ca2+ activity is required for DVE migration, we cultured E5.5 Hex-GFP embryos (where DVE cells are marked by cytoplasmic EGFP), with 10nM thapsigargin or DMSO as a negative control, and assessed the extent of their migration after 8 hours (Fig. 6A). While 71% (N=20/28) control embryos showed DVE migration (Fig. 6A-A’, D), only 10% (N=4/40) of thaspsigargin treated embryos showed DVE migration (Fig. 6C-C’, D, also see Fig. S4A-B), with

**Figure 6.**
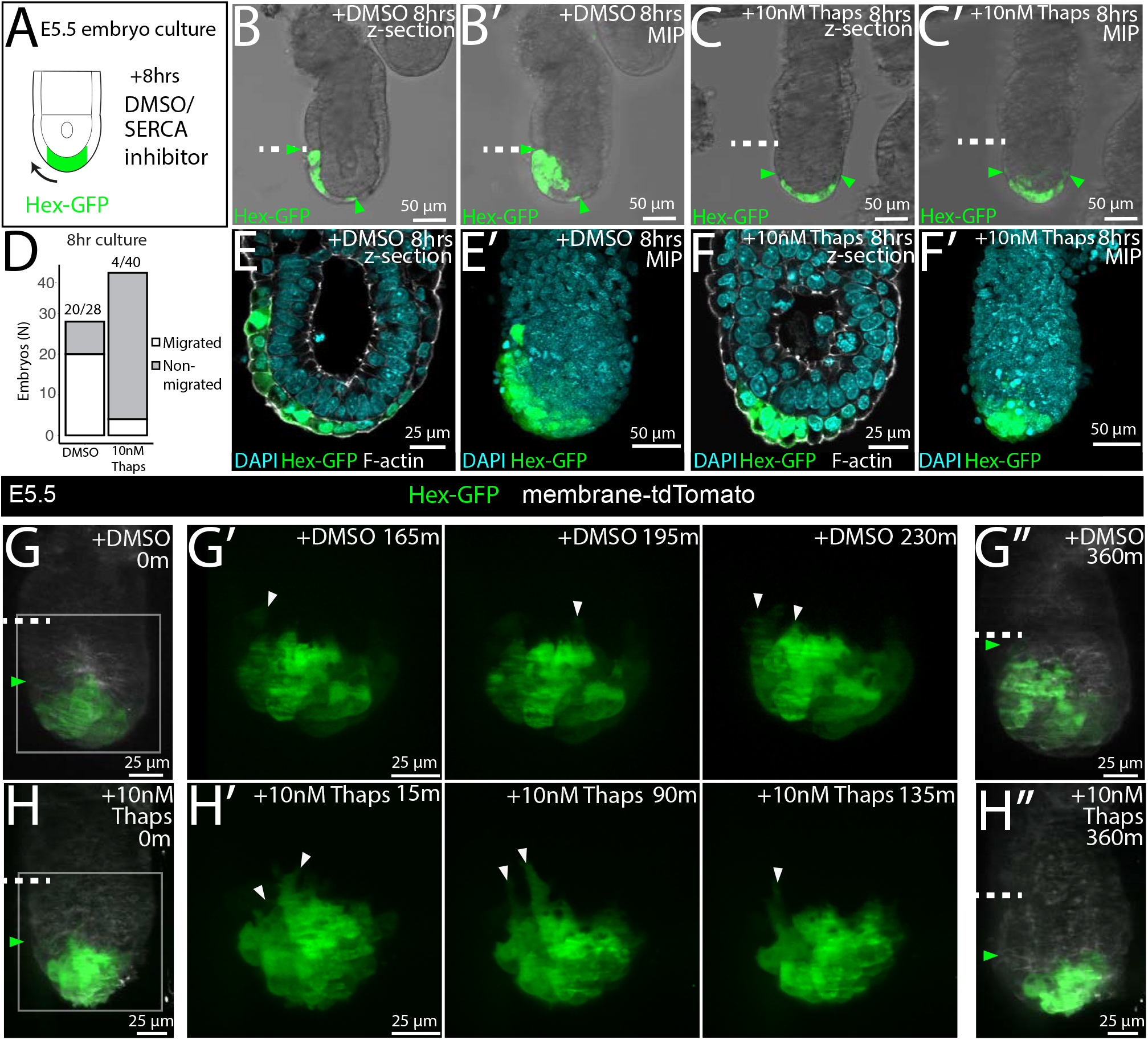
Inhibition of SERCA arrests DVE migration at E5.5, but does not block DVE cell projections. **(A)** Diagram of experimental method showing the culture of Hex-GFP embryos with 10nM thapsigargin or 0.1% DMSO or as a control. **(B-B’)** Example E5.5 embryo showing a single z-section and max-intensity projection (MIP) after 8 hours culture with DMSO (B-B’) or 10nM thapsigargin **(C-C’)** (green arrows mark position of the DVE, white dashed-line the emVE-exVE boundary). **(D)** Graph showing the number of embryos that migrated in DMSO or 10nM thapsigargin treatment after 8 hours. While 71% (20/28) of DMSO controls embryos migrated, only 10% (4/40) of thaspigargin treated embryos were able to migrate. **(E-F’)** Example Hex-GFP embryos post-fixation after 8 hour culture showing DMSO and 10nM thapsigargin treated embryos stained with DAPI and Phalloidin (F-actin) showing that VE cells remain as an intact monolayer in both control and thapsigargin treated embryos. **(G-H)** Example of E5.5 embryos imaged at 5 minute intervals for 6 hours using a Lattice Lightsheet microscope while being cultured in DMSO **(G-G’’)** or 10nM thapsigargin **(H-H’’)** (green arrowheads mark the proximal position of DVE cells, white dashed-line; emVE-exVE boundary). H’ and G’ show the Hex-GFP channel in the boxed region in H and G, for 3 selected time points during the time-lapse to highlight anteriorwards cellular projections (white arrowheads) produced in both culture conditions. G’’ and H’’ show the position of the DVE at the end of the 6 hour culture imaging period. While embryos imaged in both DMSO and thapsigargin showed cellular projections, only 7.7% (2/26) embryos culture with thapsigargin migrated in the 6 hours culture compared to 76% (13/17) of DMSO controls (see Movie S15).

Hex-GFP cells instead arrested at the distal tip of embryos (Fig. 6E-F’, Fig. S4A’). We found a similar inhibition of DVE migration when embryos were cultured in 200nM CPA (only 6/25 embryos showed DVE migration when cultured for 10 hours with 200nM CPA compared with 14/14 control embryos), suggesting that SERCA regulation of Ca2+ activity is required for DVE migration.

As Ca^2+^ signalling can also regulate a wide-range of cellular processes including proliferation, apoptosis and cell fate, we fixed embryos that had been cultured in 10nM thapsigargin or DMSO for 8 hours and performed whole-mount immunohistochemistry for key markers. We found no significant difference in the number of cell divisions as marked by Phospho-Histone H3 (Fig. S4C-C’), nor in the apoptotic marker Cleaved Caspase-3 (Fig. S4D-D’). Though there was no effect on the apoptosis or cell proliferation, thapsigargin treated embryos were on average 7% smaller than DMSO cultured controls (Fig. S4E). Finally, we found no difference in key cell identity markers of the epiblast (OCT4) (Fig. S5A) or extra-embryonic ectoderm (CDX2) (Fig. S5B), nor to the distribution of F-actin (Fig. S5C-C’).

### Cellular projections in DVE cells are not inhibited by thapsigargin treatment

We next investigated the cellular mechanism underpinning thapsigargin-mediated DVE arrest. As Ca^2+^ signalling has been shown to regulate cell polarity and filopodial projections in migrating cells (Tsai and Meyer 2012, Tsai, Kuo et al. 2015, Kim, Lee et al. 2016) and DVE cells generate basally-located projections polarised in the direction of their migration (Srinivas, Rodriguez et al. 2004, Migeotte, Omelchenko et al. 2010) we investigated whether thapsigargin blocked DVE cellular projections. We imaged E5.5 Hex-GFP embryos treated with 10nM thapsigargin or DMSO as a negative control, using high resolution lattice lightsheet microscopy to capture entire image volumes every 5 minutes over the course of 6 hours. In 76% (N=13/17) of control embryos, the DVE successfully migrated within 6 hours, and DVE cells in these embryos showed basal projections (N=17/17; Fig 6.G-G’’, Movie S15). Thapsigargin treated embryos showed a failure to migrate with only 8% (N=2/26) migrating within 6 hours, but imaging revealed that DVE cells in all embryos retained dynamic cellular projections (N=26/26; Fig. 6H-H’’, Movie S15). This indicates that SERCA inhibition of DVE migration is not due to the loss of basal projections in DVE cells.

## DISCUSSION

Ca^2+^ signalling can control a diverse range of cellular processes (Clapham 2007, Paudel, Sindelar et al. 2018, Stewart and Davis 2019), but its potential role during early mouse development has not been well characterised. Using a transgenic Ca^2+^ reporter mouse, we performed a detailed assessment of Ca^2+^ activity between E0.5 - E5.5 by live imaging. We find that Ca^2+^ transients, though initially rare events, become upregulated in a tissue specific manner as development progresses. Extra-embryonic tissues such as the E5.5 VE and ExE show increased occurrences of Ca^2+^ transients but, in contrast, the pluripotent epiblast remains relatively quiescent, with only occasional, large scale Ca^2+^ waves that spread across the tissue. We show that the Ca^2+^ transients in the VE are functionally important, since inhibition of transients by blocking SERCA leads to migration arrest of the DVE cell population that is required for correct A-P patterning.

Ca^2+^ has been show to control several aspects of cell migration in other contexts including; the front-back polarity of cells (Tsai, Kuo et al. 2015, Kim, Lee et al. 2016), the behaviour of leading cells (Hayashi, Yamamoto et al. 2018), and the regulation of lamellipodia (Tsai and Meyer 2012). However, in this study we find that DVE cells do not show any differences in cytosolic Ca^2+^ level relative to surrounding VE cells, nor a clear front-to-back gradient within cells. Furthermore, we find that DVE cells retain dynamic cellular projections when cultured with inhibitors of SERCA. This indicates that such projections are either not sufficient to drive migration, or those that are present are somehow qualitatively altered so that they no longer can facilitate DVE migration, for example, by affecting the regulation of focal adhesions (Giannone, Ronde et al. 2002, D’Souza, Lim et al. 2020). Alternatively, the role of Ca^2+^ could be non-cell autonomous – DVE cells must migrate through the emVE epithelium, with surrounding cells undergoing cell-cell rearrangements to accommodate the migrating cells (Takaoka, Yamamoto et al. 2011, Trichas, Joyce et al. 2011). As Ca^2+^ activity occurs throughout the VE, it could regulate cell-cell interactions, for example via the WNT-PCP pathway (Wallingford, Ewald et al. 2001), which has also been implicated in DVE migration (Trichas, Joyce et al. 2011). Interestingly, control embryos cultured in DMSO also show a decrease in the average number of Ca^2+^ transients, but show no detectible defect in DVE migration. The reduction in transients in the DMSO controls is an order of magnitude lower than that seen upon SERCA inhibition, which suggests that is a threshold in Ca^2+^ transient activity below which DVE migration is impaired.

In addition to the VE, we also find periodic Ca^2+^ transients in the extra-embryonic ectoderm, which serves as an important developmental signalling centre (Tam and Loebel 2007, Stower and Srinivas 2018). Ca^2+^ has been shown in other contexts to regulate cell fate as well as signalling pathways involved in embryonic development (Clapham 2007, Stewart and Davis 2019) (Paudel, Sindelar et al. 2018, Mizuno, Shiozawa et al. 2020). Though we find no loss of the key cell fate marker CDX2 in the ExE upon inhibition of Ca^2+^ transients at E5.5, it is currently unclear whether any signalling pathways are affected in this tissue.

Though periodic single-cell Ca^2+^ transients occur throughout extra-embryonic tissues at E5.5, we find that they rarely occur in the epiblast. When Ca^2+^ transients are observed in the epiblast, in contrast to those observed in extra-embryonic tissues, they span multiple cells and progress from cell to cell as a wave. The difference in Ca^2+^ transient activity between embryonic and extra-embryonic tissues is similar to that seen in other species such as *Xenopus* (Leclerc, Webb et al. 2000), Zebrafish (Gilland, Miller et al. 1999) and *Drosophila* (Markova, Senatore et al. 2015), and suggests a possible conservation in Ca^2+^ handling across species with respect to the relative quiescence of pluripotent embryonic cells. Where the epiblast shows a wave of Ca^2+^ transients, we observed in a third of cases that it was preceded by a transient in a neighbouring emVE cell, suggesting that the VE cell could be acting as a ‘trigger’ to this event. Whether this is indeed the case will be important to clarify in future experiments, particularly given VE cells are in contact with the basal aspect of epiblast cells, while transients within epiblast cells propagate in a apical to basal direction.

At pre-implantation stages, we observed numerous puncta with high levels of GCAMP6f signal. These are similar to the vesicles (MARVs) and mitochondria observed in previous studies using the Fluo-4 AM dye (Hildebrandt, Wang et al. 2019, Wang, Yasmin et al. 2022), though we do not confirm their identity in this study. We also find that such puncta are not present at later stages when the cytosolic Ca^2+^ transients begin, suggesting that there is a transition in Ca^2+^ handling as development proceeds.

In conclusion we have found a novel requirement for Ca^2+^ signalling during anterior-posterior axis formation in E5.5 the mouse embryo. Future work, to understand its mechanism of action in the DVE, and other tissues at this stage will be required using targeted approaches to affect Ca^2+^ signalling in specific cell types.

## ACKNOWLEDGMENTS

We thank: Dr Christophe Royer for help with pre-implantation stages; Helena Coker at the Oxford-ZEISS Centre of Excellence in Biomedical Imaging for assistance with the lattice light-sheet 7 microscope; the Micron Advanced Bioimaging Facility, supported by Wellcome Strategic Awards 091911/B/10/Z and 107457/Z/15/Z, for assistance the Z.1 lightsheet microscope; Pathology Services Building and Biomedical Services staff for excellent animal support; and Dr Satish Arcot Jayaram for help with mouse breeding. This work was funded by: BBSRC Research Grant BB/J00989X/1, BBSRC ALERT13 Award BB/L014750/1, BBSRC Pioneer Award BB/Y513106/1, Wellcome Strategic Award 108438/Z/15/Z and Wellcome Senior Investigator Award 103788/Z/14/Z to SS; BHF Immediate Fellowship FS/18/24/33424 to R.C.V.T. and Pump-Priming funds from the BHF Oxford Centre of Research Excellence RE/18/3/34214. FZ and XL were funded by the Ludwig Institute for Cancer Research.

## SUPPLEMENTARY DATA AND TABLES

**Figure S1.**
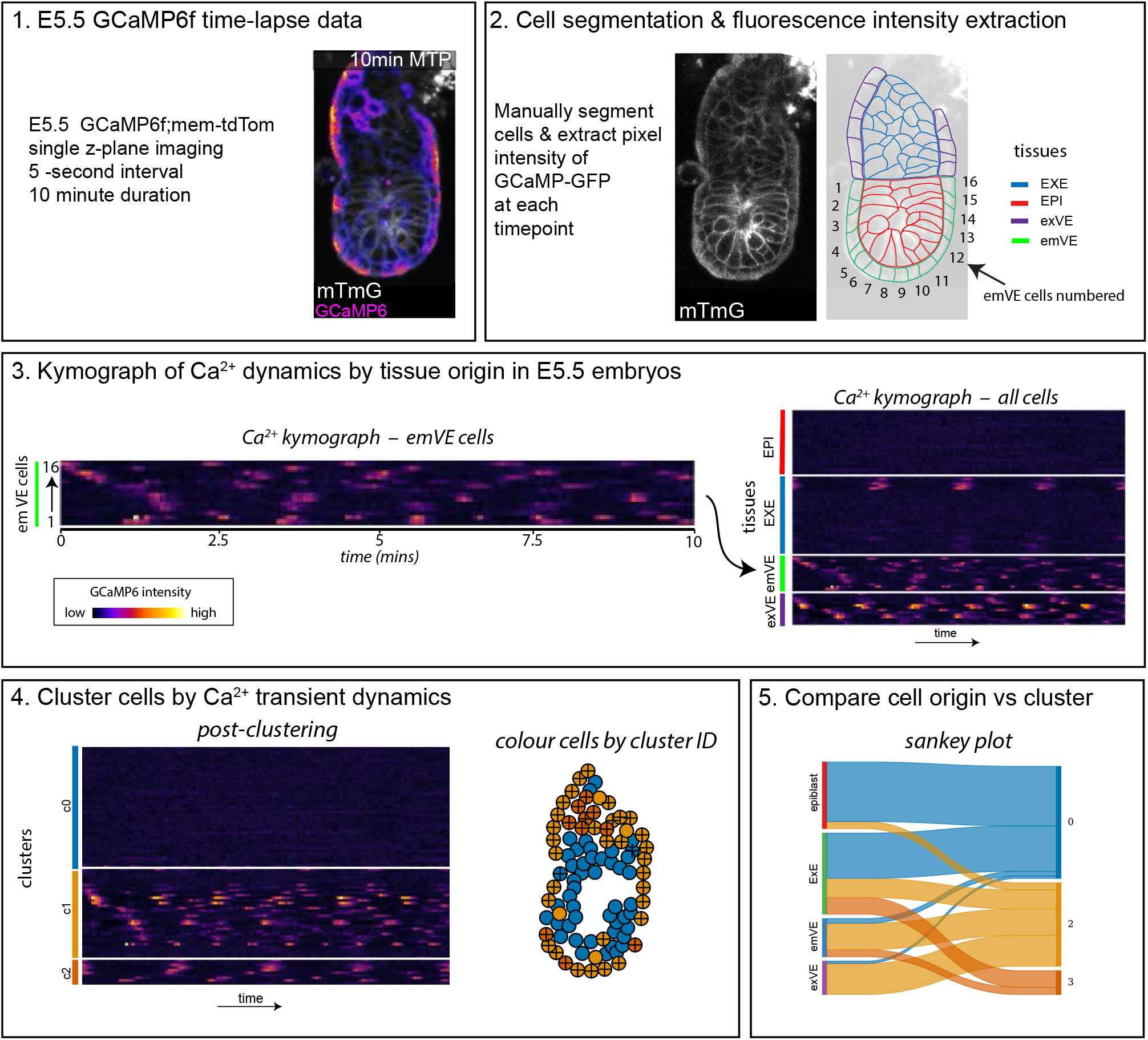
Work-flow for unbiased clustering analysis of Ca^2+^ transients at E5.5. **(1)** E5.5 GCaMP6f:membrane-tdTomato embryos (N=14) were imaged at a single z-plane for 5-second interval for 10 minutes. **(2)** Each cell was manually segmented and given an ID number. Average pixel intensity for each cell was extracted at every time-point. **(3)** GCaMP6f intensity for each embryo was plotted as a kymograph with cells ordered by tissue along the y-axis and 5-second interval timepoints along the x-axis. **(4)**. Traces of each cell were pairwise compared to construct a similarity matrix where similarity was defined as the largest positive value of the signal cross-correlation. A hierarchical density based clustering method, HDBSCAN (Campello, Moulavi et al. 2013) was then applied to automatically generate clusters. Kymographs were re-plotted to reflect the unbiased clustering and a spatial map of the embryo was coloured according to the clusters – showing here clustering results of a single embryo. **(5)** A Sankey plot was generated for each embryo to show the contribution of cells from tissues to each cluster.

**Figure S2.**
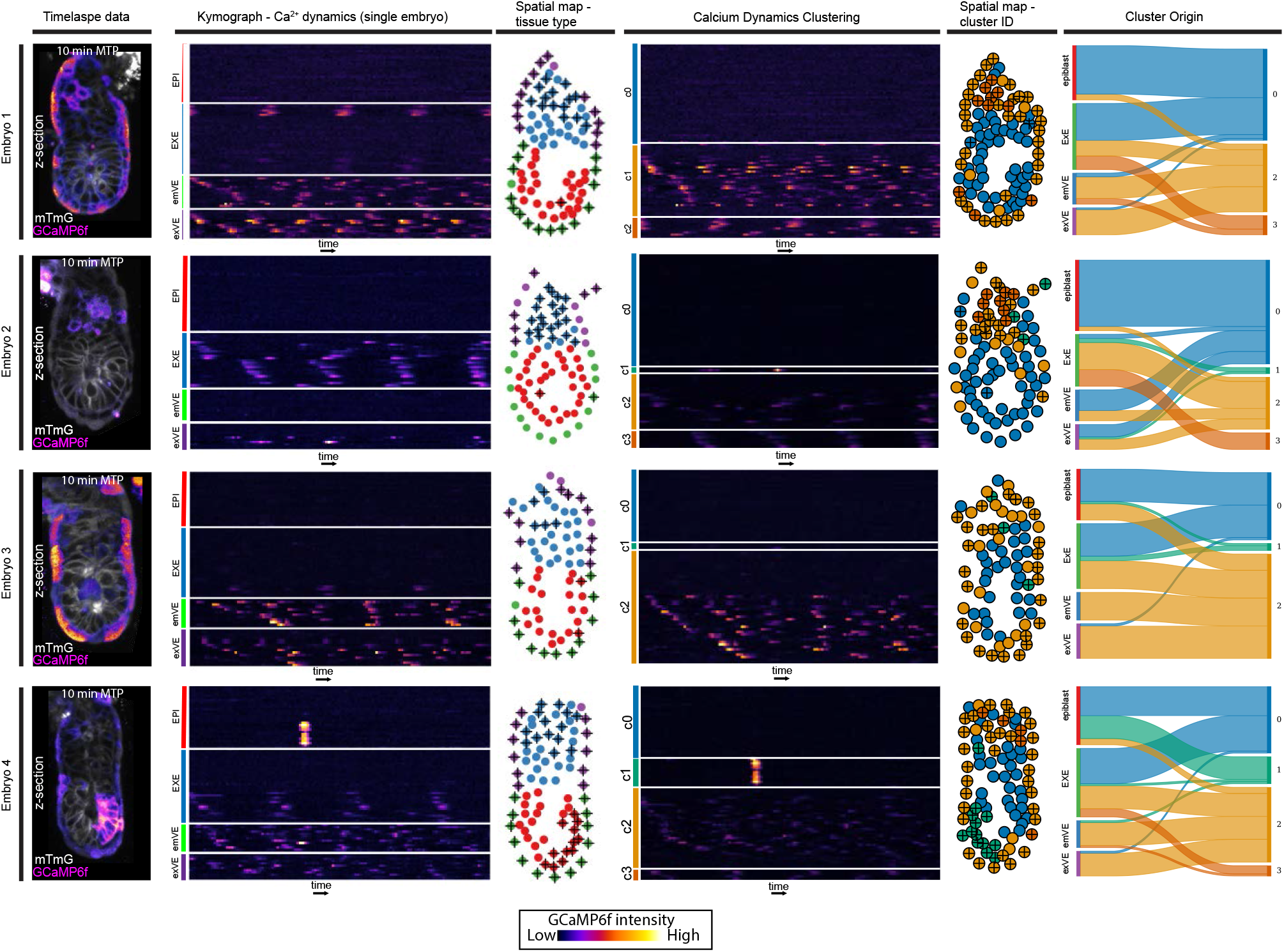
E5.5 Ca^2+^ transient clustering. E5.5 GCaMP6f:membrane-tdTomato embryos imaged imaged at a single z-plane for 5-second interval for 10 minute. First column shows max-intensity time projection (MTP) of 10 minute time-lapse. For each embryo a kymograph and spatial map are shown ordered first tissue type, then after global clustering. In the final column a Sankey graph shows the contribution of cells in each tissue to the clusters.

**Figure S3.**
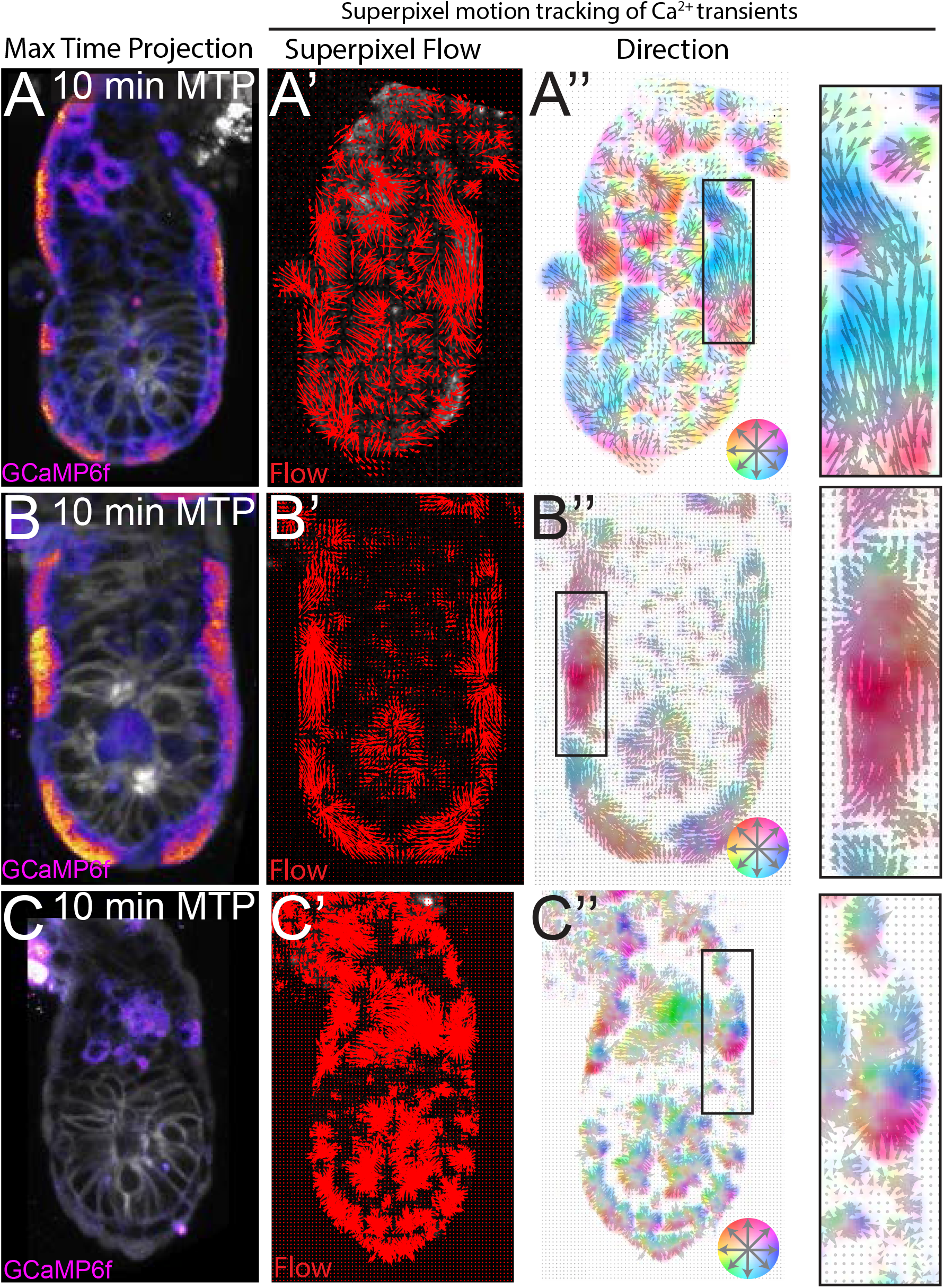
Super-pixel motion analysis shows intercellular calcium flows between adjacent cell, but no directional bias at E5.5. (A, B, C) Selected examples of E5.5 GCaMP6f:membrane-tdTomato embryos imaged every 5 seconds for 10 minutes. (A’,B’,C’) Motion Ca^2+^ transient flow using super pixel tracking. Red arrows represent the local directional motion of the superpixel during 10 min imaging duration. (A’’, B’’, C’’) Ca^2+^ transient motion tracking outputs colour-coded by vector direction, showing intercellular Ca^2+^ transients can move proximally or distally along the VE.

**Figure S4.**
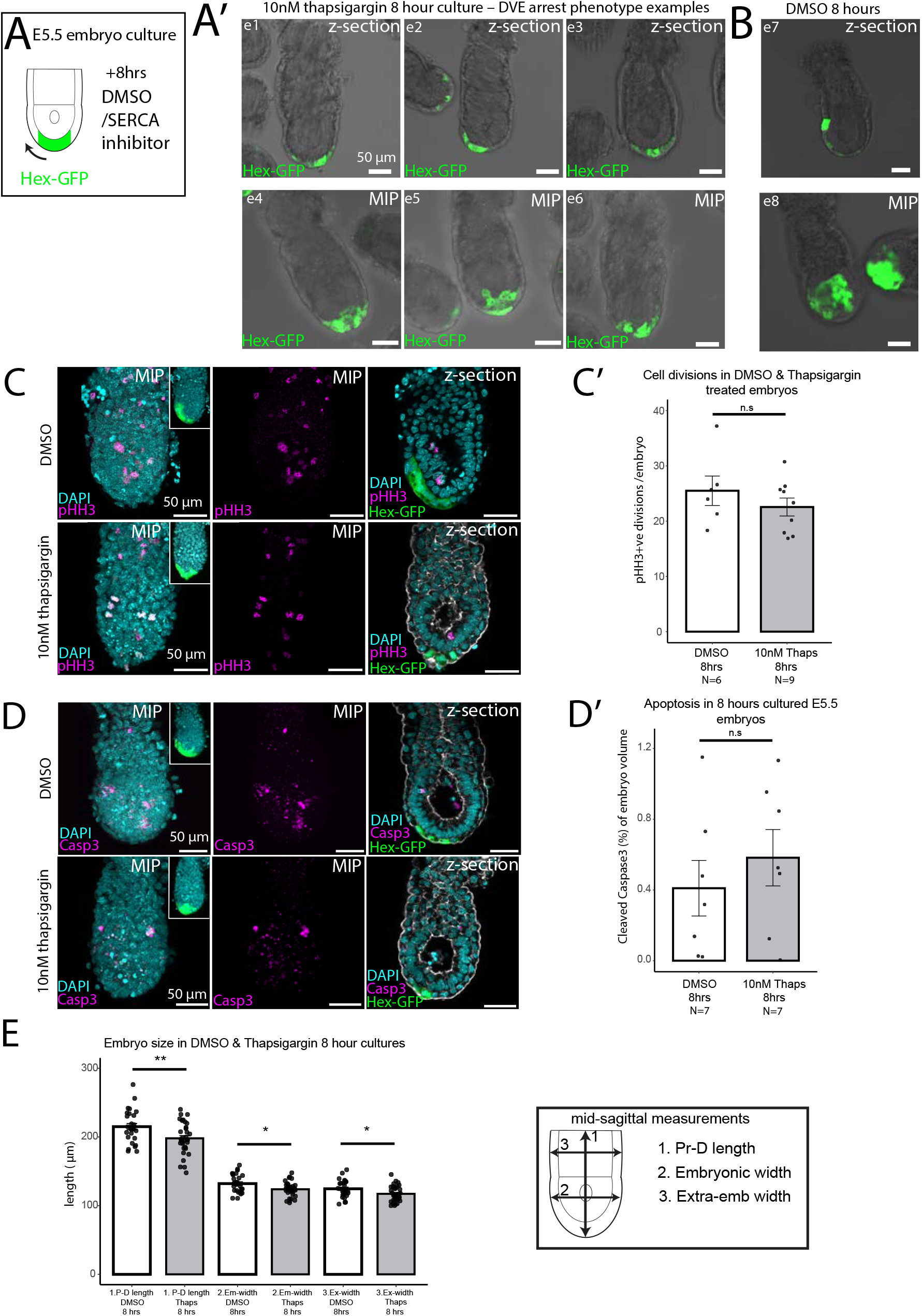
Analysis of thapsigargin cultured E5.5 embryos. **(A)** Hex-GFP embryos were cultured for 8 hours in the SERCA inhibitor thapsigargin. **(A’-A’’)** Additional examples of Hex-GFP embryos showing a single z-section (e1-e3) or max intensity projection (e4-e6) where the DVE is arrested at the distal tip after 8 hours culture with 10nM thapsigargin. **(B)** Example of DMSO control embryos successfully migrated after 8 hours. **(C-D)** Example whole-mount immunohistochemistry staining of cell proliferation marker, Phospho-HistoneH3 (pHH3) and apoptotic marker Cleaved Caspase-3 (Casp3) in cultured embryos. **(C’-D’)** Quantification of cell proliferation and cell apoptosis showed no significant different between controls (N=7) and cultured embryos (N=7) (Student’s T-test, p=>0.05). **(E)** Embryos cultured in 10nM thapsigargin (N=34) for 8 hours were slightly smaller than DMSO (N=27) control cultured embryos (Student’s T-test: Proximal-distal length p=<0.01, embryonic width p=<0.05, extra-embryonic width p=<0.05).

**Figure S5.**
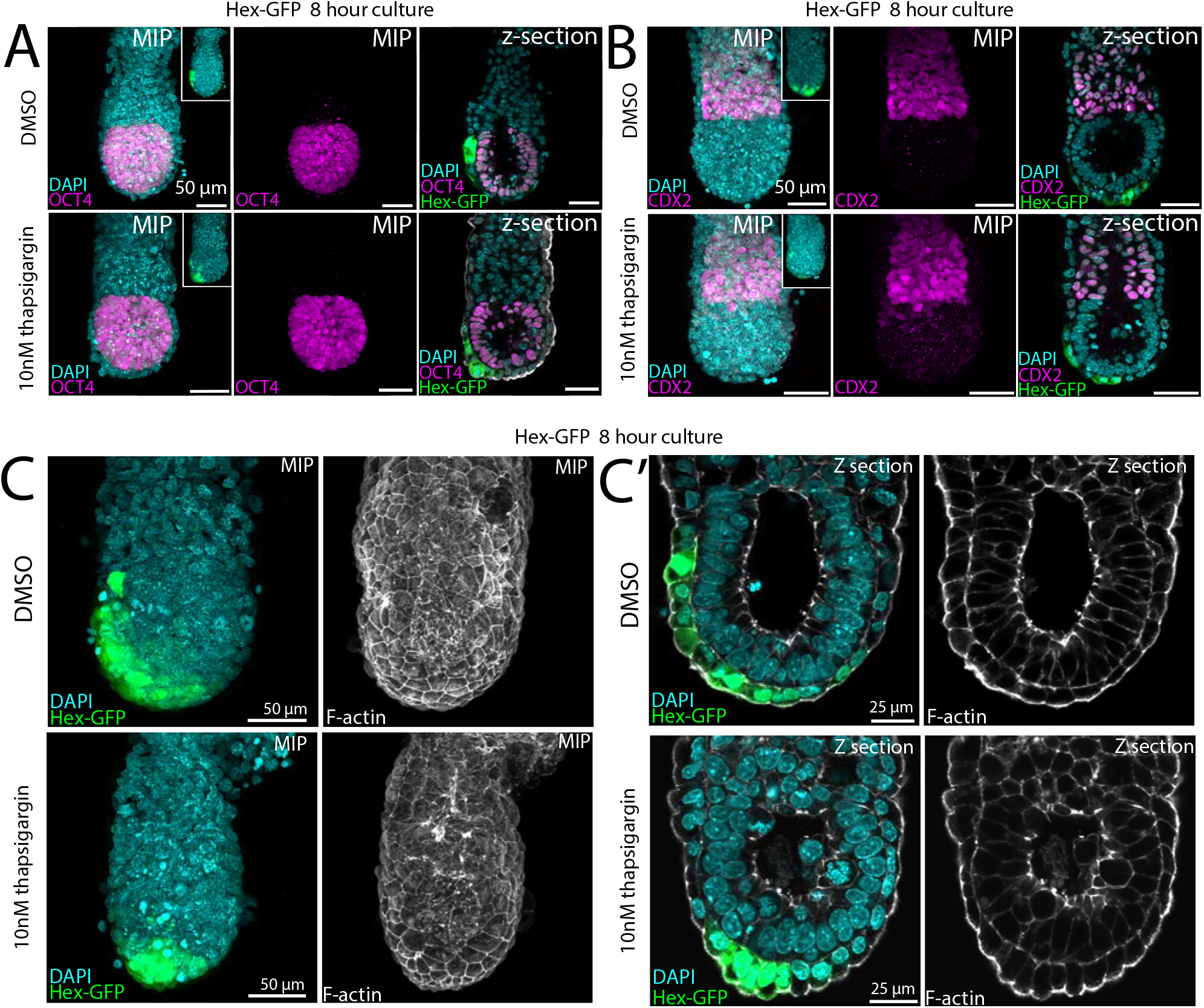
Whole-mount immunofluorescence of cell fate markers and F-actin in thapsigargin cultured E5.5 embryos. Hex-GFP embryos cultured for 8 hours in the SERCA inhibitor thapsigargin or DMSO control and stained for; **(A)** epiblast marker OCT4 (DMSO N=4, 10nM thapsigargin N=6) **(B)** extra-embryonic ectoderm markers CDX2 (DMSO N=5, 10nM thapsigargin N=4), no difference was seen in either marker. **(C)** Example embryo showing phalloidin staining on cultured embryos showing that the F-actin cytoskeleton is unaffected in thapsigargin cultured embryos (N=18) compared to DMSO (N=15) and that VE cells remain as a monolayer epithelium. Max-intensity projection (MIP).

**Table S1.**
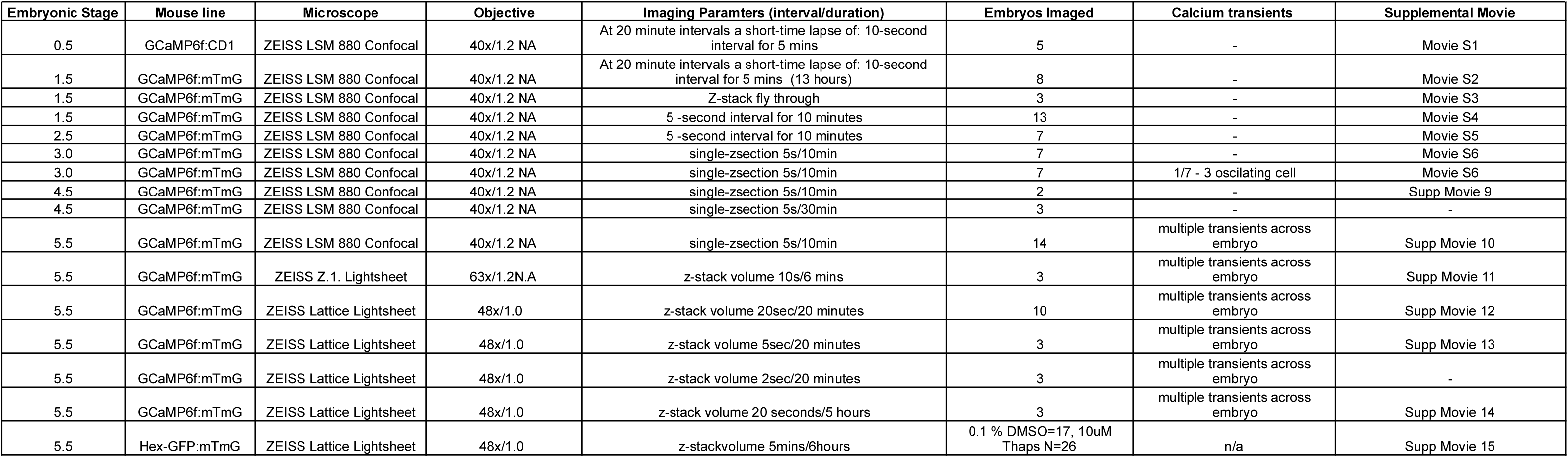

**Table S2.**

| Table S2 |  |  |  |  |
| --- | --- | --- | --- | --- |
| E5.5 tissue-based analysis of calcium transients |  |  |  |  |
|  | Epiblast | EXE | emVE | exVE |
| Total cells analysed (N=14 embryos) | 323 | 430 | 205 | 168 |
| Number of cells $\geq 1$ Ca<sup>2+</sup> transient | 32 | 163 | 83 | 94 |
| Percentage of cells $\geq 1$ Ca<sup>2+</sup> transient | | | | |
| | Percentage of cells $\geq 1$ Ca<sup>2+</sup> transient | StDev | StError | N |
| epiblast | 8.1 | 11.1 | 2.95 | 32 |
| ExE | 36.5 | 22.1 | 5.91 | 163 |
| emVE | 41.6 | 34.6 | 9.25 | 83 |
| exVE | 55.2 | 31.5 | 8.42 | 94 |
| Average number of transients per cell (in cells with $\geq 1$ Ca<sup>2+</sup> transient) | | | | |
|  | Average oscillations | StDev | StError | N |
| epiblast | 1.25 | 0.5 | 0.25 | 5 |
| ExE | 8.92 | 7.7 | 2.04 | 125 |
| emVE | 5.1 | 5 | 1.35 | 71 |
| exVE | 6.21 | 4.1 | 1.09 | 87 |
| Periodicity of Calcium oscillations |  |  |  |  |
|  | Average Periodicity (min:sec) | StDev | StError | Oscillations (N) |
| epiblast | 1:54 | 1.45 | 0.65 | 5 |
| ExE | 3:00 | 1.41 | 0.13 | 125 |
| emVE | 2:32 | 0.95 | 0.11 | 71 |
| exVE | 2:54 | 1.27 | 36462760654 | 87 |
| Average peak transient Intensity |  |  |  |  |
|  | Average intensity | StDev | StError | N |
| epiblast | 33.29 | 38.09 | 7.2 | 32 |
| ExE | 8.45 | 6.16 | 0.48 | 163 |
| emVE | 13.61 | 10.22 | 1.12 | 83 |
| exVE | 14.25 | 14.25 | 1.1 | 94 |
| Average peak transient duration |  |  |  |  |
|  | Average duration (sec) | StDev | StError | N |
| epiblast | 23.96 | 20.08 | 3.79 | 32 |
| ExE | 21.15 | 8.95 | 0.72 | 163 |
| emVE | 18.93 | 11.62 | 1.28 | 83 |
| exVE | 14.39 | 6.98 | 0.69 | 94 |

**Table S3.**

| Table S3 |  |  |  |  |
| --- | --- | --- | --- | --- |
| E5.5 cluster-based analysis of calcium transients |  |  |  |  |
|  | Epiblast | ExE | emVE | exVE |
| Total cells analysed (N=14 embryos) | 323 | 430 | 205 | 168 |
| Cluster 0 | 272 | 204 | 96 | 61 |
| Cluster 1 | 14 | 12 | 7 | 10 |
| Cluster 2 | 37 | 148 | 97 | 109 |
| Cluster 3 | 0 | 49 | 5 | 5 |
| Average number of transients per cell (in cells with $\geq 1$ Ca<sup>2+</sup> transient) | | | | |
|  | Average oscillations | StDev | StError |  |
| Cluster 0 | 0.02 | 0.2 | 0.010 |  |
| Cluster 1 | 1.51 | 0.8 | 0.126 |  |
| Cluster 2 | 2.21 | 2.0 | 0.103 |  |
| Cluster 3 | 3.80 | 1.3 | 0.175 |  |
| Periodicity of calcium oscillations |  |  |  |  |
|  | Average Periodicity (min:sec) | StDev | StError |  |
| Cluster 0 | 3:00 | 1.45 | 1.63 |  |
| Cluster 1 | 3:27 | 1.41 | 0.33 |  |
| Cluster 2 | 2:40 | 0.95 | 0.10 |  |
| Cluster 3 | 2:21 | 1.27 | 0.11 |  |
| Average peak transient Intensity |  |  |  |  |
|  | Average Intensity (AU) | StDev | StError |  |
| Cluster 0 | 3.9 | 2.1 | 0.75 |  |
| Cluster 1 | 30.0 | 31.8 | 4.85 |  |
| Cluster 2 | 10.7 | 8.6 | 0.53 |  |
| Cluster 3 | 11.9 | 7.7 | 1.04 |  |
| Average peak transient duration |  |  |  |  |
|  | Average Duration (sec) | StDev | StError |  |
| Cluster 0 | 29.19 | 28.18 | 6.46 |  |
| Cluster 1 | 15.08 | 5.60 | 0.85 |  |
| Cluster 2 | 18.74 | 11.37 | 0.70 |  |
| Cluster 3 | 19.24 | 4.28 | 0.57 |  |

